# Transient suppression of the ECM1 gatekeeper is essential for HGF/c-MET-driven liver regeneration

**DOI:** 10.64898/2026.06.08.730887

**Authors:** Ye Yao, Yujia Li, Chenjun Huang, Jiayi Ma, Liang Xu, Stephanie D. Wolf, Julian Gilljam, Haifeng Zeng, Stefan Munker, Xioachun Cao-Ehlker, Ruochan Li, Bernhard Renz, Hanno Nieß, Seddik Hammad, Elisa Holstein, Laura Danielczyk, Roman Liebe, Honglei Weng, Ursula Klingmüller, Matthias P. A. Ebert, Chunfang Gao, Johannes Bode, Donato Inverso, Peter ten Dijke, Shanshan Wang, Stefan Höhme, Weiguo Fan, Steven Dooley, Sai Wang

## Abstract

Hepatocyte growth factor (HGF) is a multifunctional cytokine stored in the extracellular matrix as an inactive precursor and is essential for tissue repair. How HGF activity is dynamically regulated during regeneration remains unclear. Here, we identify extracellular matrix protein 1 (ECM1) as a physiological inhibitor of active HGF during liver regeneration. Following 70% partial hepatectomy, active HGF rapidly increases in parallel with a sharp decline in ECM1, and preventing this downregulation delays liver mass recovery. Mechanistically, ECM1 directly binds the active HGF α-subunit through a mechanism dependent on residue R392, thereby suppressing c-MET–ERK–MYC signaling and hepatocyte proliferation. Loss of ECM1 permits activation of this pathway, whereas MYC overexpression rescues ECM1-mediated growth inhibition. In patients, proliferative hepatocytes localize to ECM1-negative regions. Supported by transcriptomic analyses and computational modeling, these findings identify ECM1 as an extracellular gatekeeper whose transient downregulation enables HGF-driven tissue repair.

**Highlights:**

- ECM1 is temporally downregulated during liver regeneration, while forced ECM1 expression delays liver mass recovery after 70% PHx by suppressing hepatocyte proliferation.
- Rapid and transient ECM1 downregulation permits HGF-c-Met-ERK-MYC signaling to drive hepatocyte cell-cycle entry and expansion.
- ECM1 directly binds the active HGF α-subunit with residue R392 playing a critical role, revealing a novel extracellular mechanism for growth factor inhibition.
- Loss of ECM1 marks proliferative hepatocytes in human liver repair, identifying ECM1 as a tunable regulator of regenerative capacity.

## Introduction

Hepatocyte growth factor (HGF) is a multifunctional cytokine that regulates epithelial cell growth, morphogenesis, and tissue repair across a variety of organs (Fausto, 2000; Michalopoulos, 2010). The liver has a remarkable capacity to regenerate after injury, restoring its mass and function through tightly coordinated cellular and molecular programs (Michalopoulos, 2007; Michalopoulos and Bhushan, 2021). A large amount of HGF is stored in the extracellular matrix of healthy liver in its precursor (pro-HGF) latent form, which can be activated via proteolytic cleavage at Arg494–Val495 residues by the HGF activator (HGFA), urokinase-type plasminogen activator (uPA), plasma kallikrein, and metalloproteinases, yielding the biologically active α and β subunits (Mizuno et al., 1994; Naka et al., 1992; Roselli et al., 1998). Cleaved HGF uniquely stimulates its tyrosine kinase receptor c-MET, triggering intracellular JAK/STAT3, PI3K/AKT, NFκB, and RAS/ERK signaling pathways to drive hepatocyte proliferation (Fausto, 2000; Fausto et al., 1995; Michalopoulos, 2011). After a 70% partial hepatectomy (PHx), HGF activation increases within 3-72 h and returns to baseline 7-8 days post-PHx (Fausto, 2000; Karkampouna et al., 2016; Karkampouna et al., 2012; Michalopoulos and Bhushan, 2021). However, the dynamic regulation of the active form of HGF during liver regeneration remains largely uncharacterized.

Our previous studies revealed that extracellular matrix protein 1 (ECM1) is essential for liver homeostasis, which is sustained by HGF and epidermal growth factor under healthy conditions (Fan et al., 2019; Li et al., 2025). Upon liver injury, an interferon (IFN)-γ/nuclear factor erythroid 2-related factor 2 (NRF2) axis becomes activated in the stressed hepatocytes, disrupts growth factor-relevant signaling pathways, and reduces ECM1 expression, unleashing active transforming growth factor-Β (TGF-β) to drive fibrogenesis (Li et al., 2025; Link et al., 2025). It is unknown, and therefore the subject matter of the present investigation, whether HGF, the main driver for liver regeneration, and ECM1 interact during the regenerative response to liver injury. We investigated the expression dynamics of ECM1 and its functional consequences in a 70% PHx mouse model. We further examined the physical interaction between ECM1 and HGF and its impact on proliferative signaling in primary mouse hepatocytes, the human hepatocyte cell line HepaRG, and liver tissues from patients with cirrhosis or acute liver failure. In addition, we developed a detailed three-dimensional spatio-temporal multiscale (3D-STM) model that validates our experimental findings, supported the mechanistic conclusions, and enabled prediction of dynamic interactions during liver regeneration.

## Results

### Preventing Ecm1 downregulation delays liver regeneration

Given that most of the mouse liver mass is restored within 7-8 days after 70% partial hepatectomy (PHx) (Fausto, 2000; Michalopoulos and Bhushan, 2021), we monitored Hgf signaling activation and *Ecm1* expression in *Ecm1*-tdTomato mice at 2 h and days 1, 2, 4, and 8 post-PHx (**Figure S1A, B**). Consistent with previous findings (Michalopoulos, 1993), Hgf signaling was rapidly induced, detectable as early as 2 h, peaked at day 2, and returned to basal levels by day 8, as evidenced by c-Met upregulation and phosphorylation (**Figure S1C**). Unexpectedly, Ecm1 expression was markedly downregulated during liver regeneration (LR), as shown at both mRNA and protein levels, as well as by the loss of red fluorescent signal in *Ecm1*-tdTomato mice (**Figure 1A-E**). Since our previous study showed that damage stress-induced IFN-γ/ NRF2 signaling may downregulate ECM1 expression in hepatocytes (Li et al., 2025), we examined mRNA expression of the IFN-γ signaling-related genes in RNA-Seq data from mice subjected to 70% PHx. We observed an upregulated expression of IFNγ receptors (*Ifngr1* and *Ifngr2*), *Nrf2* (*Nfe2l2*), and its target genes *Cyp1b1* and *Nqo1* at day 2 post-surgery (**Figure S1D**), suggesting that IFNγ/NRF2 may contribute to the rapid ECM1 downregulation upon acute injury, although this requires further validation.

**Figure 1.**
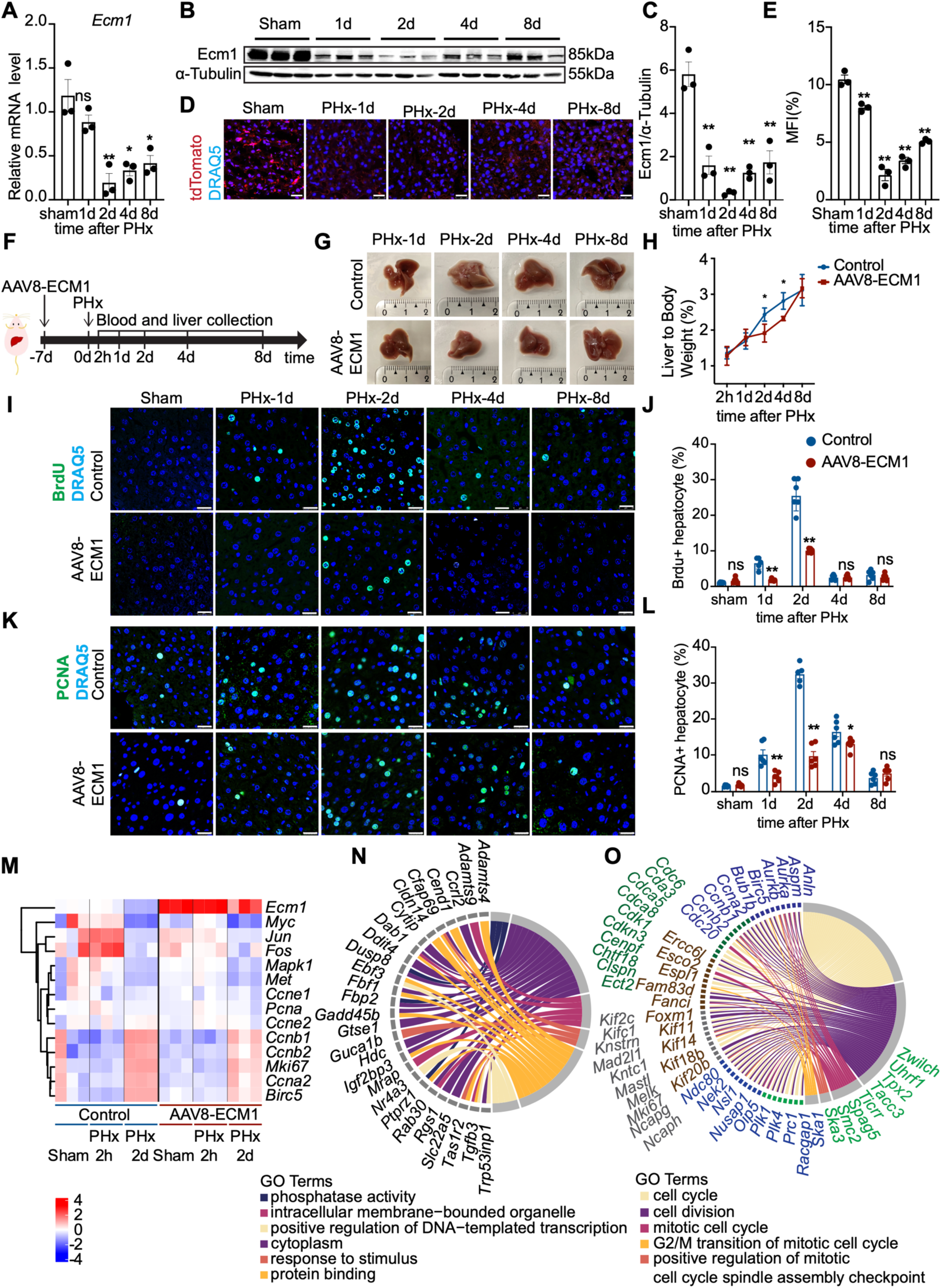
Preventing Ecm1 downregulation by AAV8-ECM1 delays liver regeneration. **(A, B)** Relative *Ecm1* mRNA and Ecm1 protein expression in liver tissues collected from sham or 70% PHx mice at days 1, 2, 4, and 8 (n=6 per group). **(C)** Quantification of Ecm1 protein levels from (B). **(D, E)** Fluorescent signal of *Ecm1*-tdTomato in liver tissues at the indicated time points after 70% PHx. **(F)** Experimental scheme showing AAV8-ECM1 administration and sample collection timeline after 70% PHx. **(G, H)** Representative liver images and liver-to-body weight ratios from sham or 70% PHx mice with control or AAV8-ECM1 treatment. **(I-L)** Representative images and quantification of IF staining for BrdU and PCNA in liver tissues from mice injected with control or AAV8-ECM1, followed by sham or 70% PHx. **(M)** RNA-seq analysis of growth factor-related signaling pathways in liver tissues collected at 2 h or 2 d post sham or 70% PHx. **(N, O)** Chord diagrams showing major biological processes suppressed by AAV8-ECM1 at 2 h or 2 d after 70% PHx. For qRT-PCR, mouse *Ppia* served as an endogenous control. For immunoblotting, α-Tubulin served as a loading control. For IF staining, nuclei were counterstained with DRAQ5. Scale bar, 25 µm. Data are presented as mean ± SD. P values were calculated using an unpaired Student’s t-test. *p<0.05; **p<0.01. Quantification of immunoblots and staining was performed using ImageJ (NIH).

To directly assess the role of ECM1 in LR, we prevented Ecm1 downregulation by ectopically expressing ECM1 in the liver via an intravenously delivered recombinant adeno-associated virus construct, AAV8-ECM1, one week before PHx surgery. Liver and blood samples were collected at defined time points, as indicated (**Figure 1F**). As expected, the liver-to-body weight ratio steadily increased and nearly recovered by day 8 post-surgery in controls. However, compensation of the rapid ECM1 expression decrease by way of its heterotopic expression (**Figure S1E**) strongly delayed liver mass recovery at days 2 and 4, although regeneration eventually caught up by day 8 (**Figure 1G, H**).

### Ecm1 overexpression inhibits hepatocyte proliferation after 70% PHx

Gain of liver mass upon surgery, including PHx in humans and mice, is mainly based on de novo generation of hepatocytes. To assess hepatocyte proliferation, we analyzed liver tissues from control and AAV8-ECM1-treated mice subjected to sham operation or 70% PHx at the time points as above. Bromodeoxyuridine (BrdU) was injected 2 h prior to tissue collection. In controls, the number of BrdU-positive hepatocytes increases after PHx, peaks at day 2 during regeneration, and returns to baseline by day 8, relative to sham mice. In contrast, AAV8-ECM1-treated mice exhibit markedly fewer BrdU-positive hepatocytes, particularly at days 1 and 2 post-PHx (**Figure 1I, J**; representative whole-slide immunofluorescence is presented in **Figure S2**). Expression of the endogenous proliferation marker proliferating cell nuclear antigen (PCNA) followed the same trend (**Figure 1K, L**; representative whole-slide staining is presented in **Figure S2**). These findings indicate that preventing Ecm1 downregulation by its ectopic re-expression suppresses hepatocyte proliferation and delays liver regeneration after PHx-mediated acute liver injury.

To investigate the molecular basis of this phenotype, we performed a comparative RNA-Seq analysis on liver tissue collected at sham, 2 h, and 2 days post-PHx with control or AAV8-ECM1 injection. As expected, classical early-response genes required for LR initiation, including *Myc*, *Fos*, and *Jun*, are induced at 2 h, while Hgf target genes, key cell-cycle inducers, and proliferation markers, such as *c-Met*, *Mapk1*, *Birc5*, *Ccna2*, *Ccnb1*, *Ccnb2*, *Ki67*, and *Pcna*, are induced at day 2, all of which are notably reduced in tissue with Ecm1 overexpression (**Figure 1M**). Gene Ontology (GO) analysis (Gene Ontology, 2021) revealed that the top differentially expressed genes altered by AAV8-ECM1 at 2 h are enriched for phosphatase activity and protein-binding categories (**Figure 1N**), whereas those at day 2 are predominantly associated with cell-cycle regulation (**Figure 1O**).

Together, these data demonstrate that Ecm1 downregulation is required for the early proliferative response in liver regeneration following 70% PHx.

### AAV8-ECM1 livers exhibit reduced Hgf-c-Met-Erk-Myc activation during LR after PHx

To determine whether Ecm1 overexpression delays LR by interfering with Hgf signaling, we assessed Hgf expression, activation, and downstream effectors in control and AAV8-ECM1–treated mice subjected to sham or PHx operations. Hepatic *Hgf* mRNA levels increase from day 1 to day 4 and return to baseline by day 8 post 70% PHx, with no difference between control and AAV8-ECM1 groups (**Figure 2A**). At the protein level, Hgf increases from 2 h to 2 days post-surgery in control mice, but is suppressed in AAV8-ECM1 livers, as determined by an enzyme-linked immunosorbent assay (ELISA) that is able to detect both the precursor form as well as active HGF (**Figure 2B**; standard curve in **Figure S3A**), suggesting ECM1 is regulating HGF protein stability rather than de novo HGF synthesis. Since Hgf signaling plays an essential role during the early phase of LR (Fausto, 2000), we next examined c-Met receptor activation. In controls, c-Met phosphorylation is robust at 2 h, 1 day, and 2 days post-PHx, whereas activation is significantly blunted in AAV8-ECM1 livers, as confirmed by immunoblotting and immunofluorescence of phosphorylated (p)-c-Met, indicative of active c-Met (**Figure 2C, D**). These results indicate that ectopically expressed Ecm1 affects LR via pre-existing HGF protein stored in the extracellular matrix rather than newly synthesized HGF, the latter being more important for replenishing the consumed HGF stock (Michalopoulos, 2010).

**Figure 2.**
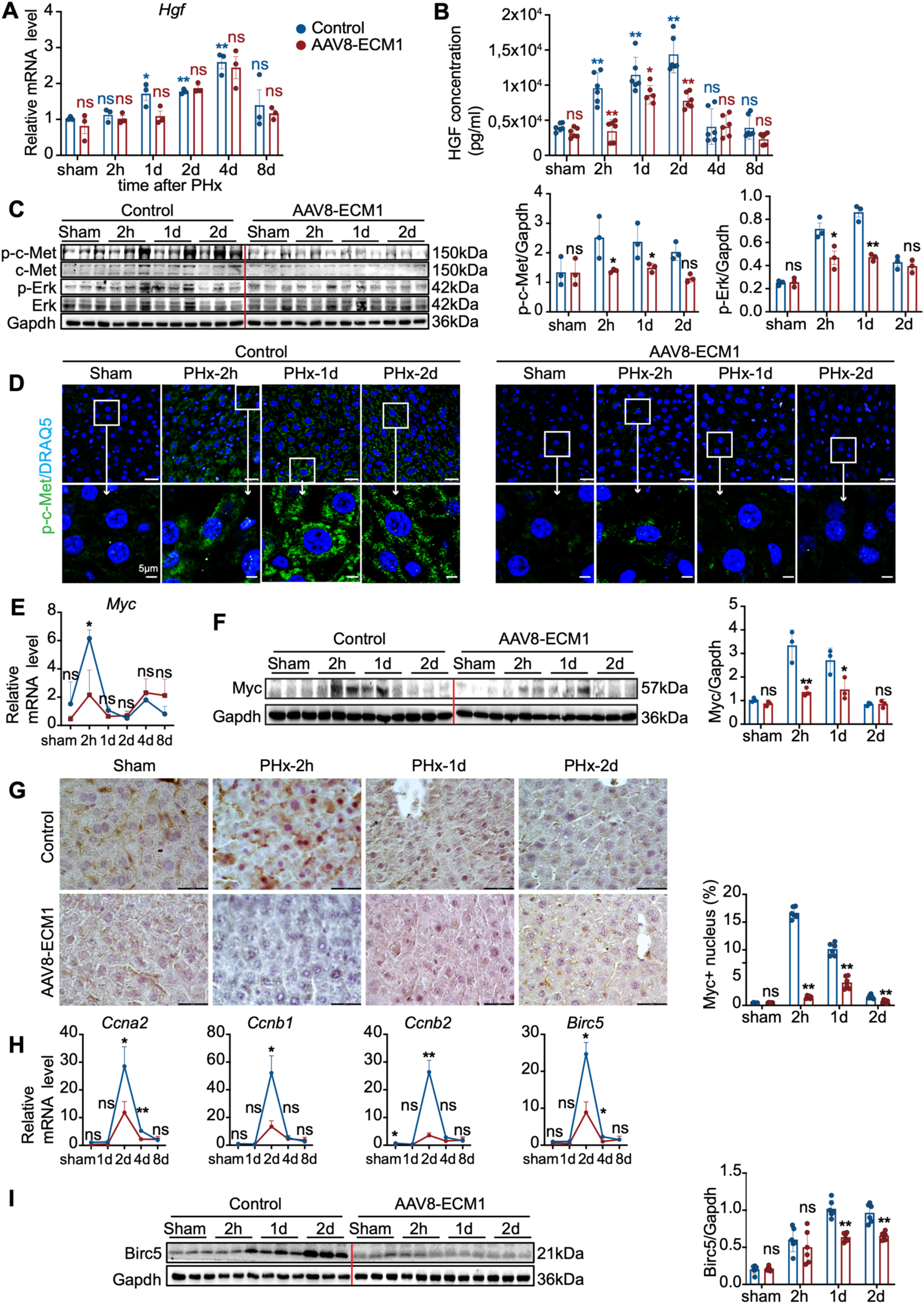
Ecm1 overexpression inhibits HGF-c-Met-Erk-Myc activation during LR. **(A)** Relative mRNA expression of *Hgf* in liver tissues from sham or 70% PHx mice collected at 2 h, 1 d, 2 d, 4 d, or 8 d. **(B)** ELISA quantification of total HGF (precursor and active forms) in the same samples. **(C)** Immunoblotting analysis of phosphorylated c-Met and Erk in liver tissues collected at 2 h, 1 d, or 2 d after sham or 70% PHx. **(D)** IF staining for p-c-Met in liver tissues at the indicated time points. Nuclei were counterstained with DRAQ5. Scale bar, 25 µm. **(E-G)** Relative mRNA expression, protein levels, and IHC staining of Myc in livers at 2 h, 1 d, or 2 d post PHx. Scale bar, 43.5 µm. **(H, I)** Relative mRNA and protein expression of cell-cycle regulators (Birc5, Ccna2, Ccnb1, Ccnb2) in livers collected at 2 h, 1 d, or 2 d post 70% PHx. For qRT-PCR, mouse *Ppia* was used as an endogenous control. For immunoblotting, α-Tubulin or Gapdh served as loading controls. Data are presented as mean ± SD. P values were calculated using an unpaired Student’s t-test. *p<0.05; **p<0.01. Quantification of staining and immunoblotting was performed using ImageJ (NIH).

We then analyzed downstream effectors. Myc is a known HGF target in the context of hepatocyte proliferation and Myc expression at both mRNA and protein levels as well as its nuclear translocation, are rapidly induced after PHx, as shown by qRT-PCR, immunoblotting, and immunohistochemistry (IHC) (**Figure 2E-G**). Notably, these responses are prevented by Ecm1 overexpression (**Figure 2E-G**). Importantly, upregulation of Myc target genes (*Birc5*, *Ccna2*, *Ccnb1*, and *Cccnb2*) that is critical for cell-cycle progression at day 2 post-PHx, is markedly suppressed in AAV8-ECM1 livers (**Figure 2H, I**). We also examined other HGF/c-Met downstream targets relevant to LR initiation, including Fos/Jun, Akt, and Stat3. *Fos/Jun* mRNA induction at 2 h post-PHx is reduced in AAV8-ECM1 livers, although protein levels remain unchanged (**Figure S3B, C**). Phosphorylations of Stat3 and Akt are not affected by Ecm1 overexpression (**Figure S3D**).

To test whether ECM1 interferes with HGF activation at the proteolytic level, we measured the levels of proteases known for HGF precursor cleavage, including urokinase plasminogen activator (*Plau/uPA*), its receptor (*Plaur*), plasma kallikrein (*Klkb1*), and matrix metalloproteinases (*Mmp2*, *Mmp9*), with uPA being particularly important during LR (Fajardo-Puerta et al., 2016; Roselli et al., 1998). Although *Plau* mRNA expression is not elevated, consistent with its regulation primarily at the activity level, expression of its receptor, *Plaur,* is strongly upregulated at 2 h post-PHx in both the control and AAV8-ECM1 groups, with no obvious difference between the two conditions (**Figure S3E**). Expression of *Klb1*, *Mmp2*, and *Mmp9* are not significantly altered by PHx or AAV-ECM1 treatment (**Figure S3F**). These findings suggest that ECM1 does not affect pro-HGF cleavage, but may instead interfere with the generation or availability of the active HGF.

### Ecm1 overexpression inhibits active HGF-induced hepatocyte proliferation *in vitro*

To test whether ECM1 directly suppresses HGF signaling, we overexpressed Ecm1 by a plasmid expression vector (vEcm1) in primary mouse hepatocytes (PMHs), and stimulated them with recombinant active HGF. The pcDNA3.1 plasmid was used as a vector control (vCon). Within 10 min, HGF induces robust c-Met and Erk phosphorylation, which is markedly inhibited by vEcm1, as shown by immunoblotting and IF staining (**Figure 3A, B**). Similar ECM1-mediated signaling inhibition is present following 24-48 h of HGF treatment (**Figure 3C, D**). Consistently, HGF-induced Myc expression is also suppressed by vEcm1 (**Figure 3C, D**). Since we observed that Myc nuclear translocation peaks at 4 h after HGF stimulation, we treated PMHs with HGF for 4 h following Ecm1 overexpression. vEcm1 strongly impairs HGF-induced nuclear accumulation of Myc (**Figure 3E**). Moreover, HGF treatment for 24 or 48 h induces Myc and cell-cycle regulator expression (Birc5, Ccna2, Ccnb2), all of which are suppressed by Ecm1 overexpression (**Figure 3F, G**). Functionally, Ecm1 overexpression inhibits HGF-driven PMH proliferation, as determined by dynamic IncuCyte life cell monitoring and a Cell Counting Kit (CCK)-8 assay (**Figure 3H, I**). To extend these findings, we performed colony-formation assays using the murine hepatocyte line AML12. HGF treatment significantly enhances colony formation, whereas ECM1 overexpression abrogates this effect (**Figure 3J**). Further, ectopic Myc expression induces expression of cell-cycle genes at mRNA and protein level, even in the presence of Ecm1 overexpression (**Figure 3K, L**). However, Myc overexpression alone is insufficient to induce primary hepatocyte proliferation *in vitro* (**Figure 3M**).

**Figure 3.**
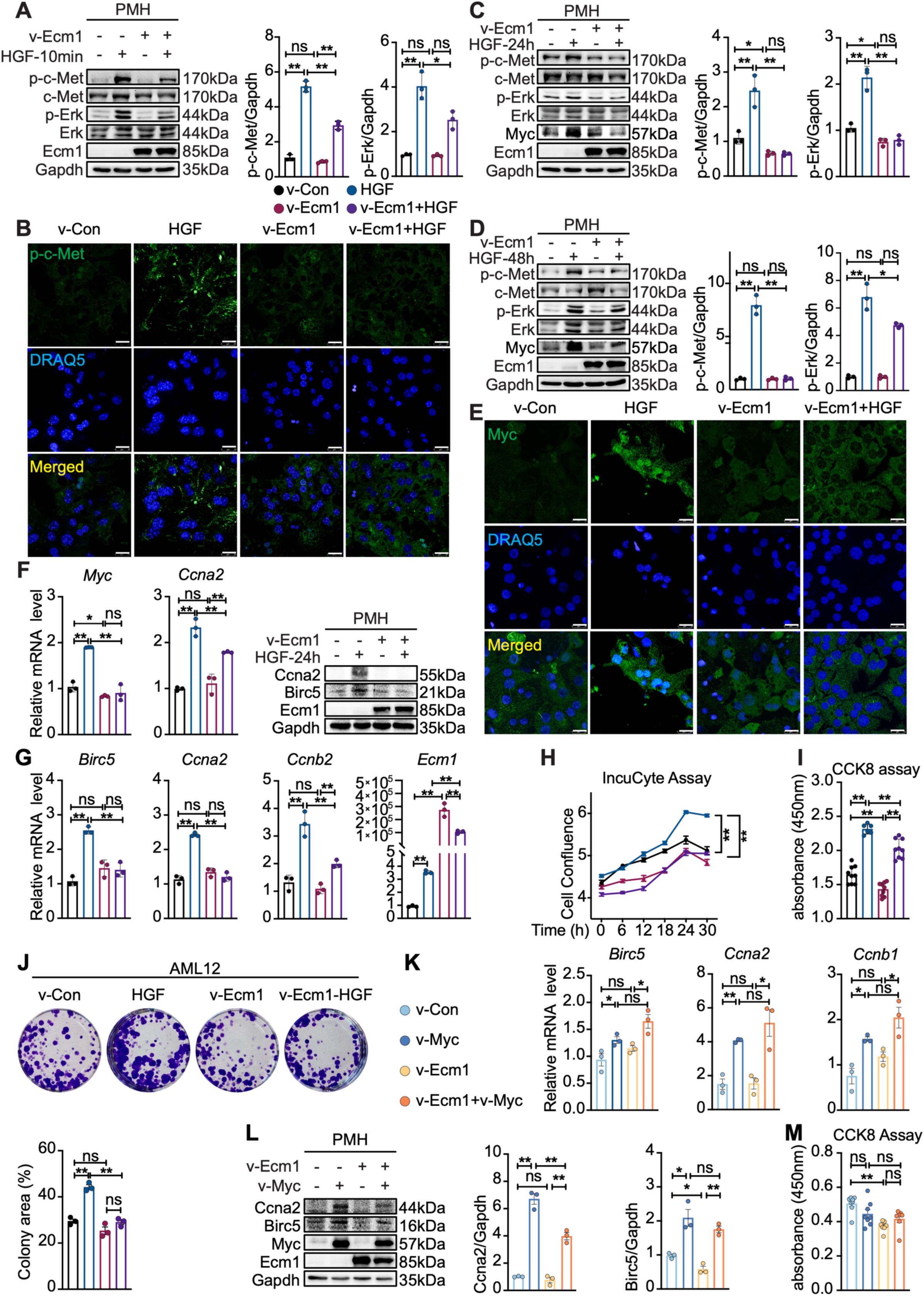
Ecm1 overexpression inhibits active HGF-induced cell proliferation in PMHs. **(A, B)** Immunoblotting and IF staining of p-c-Met and p-Erk in PMHs transfected with control or Ecm1 plasmid and stimulated with recombinant HGF for 10 min. **(C, D)** Immunoblotting analysis of p-c-Met, p-Erk, and Myc in PMHs transfected with control or Ecm1 plasmid (v-Ecm1) and treated with HGF for 24 h or 48 h. **(E)** IF staining of Myc in PMHs transfected with control or Ecm1 plasmid and treated with HGF for 4 h. **(F, G)** Relative mRNA and protein expression of cell-cycle regulators (Birc5, Ccna2, Ccnb1, Ccnb2) in PMHs transfected with control or Ecm1 plasmid and treated with HGF for 24 or 48 h. **(H, I)** Proliferation of PMHs transfected with control or Ecm1 plasmid and treated with HGF for 48-72 h, quantified by IncuCyte live-cell imaging and cell counting kit (CCK)-8 assay. **(J)** Colony formation of AML12 cells transfected with control or Ecm1 plasmid and cultured with or without HGF for 2 weeks (HGF added every 2 d). **(K, L)** Relative mRNA and protein expression of cell-cycle genes in PMHs transfected with Myc, Ecm1, or both. **(M)** Proliferation of PMHs overexpressing Myc and/or Ecm1 by CCK-8 assay. For qRT-PCR, mouse *Ppia* served as an endogenous control. For immunoblotting, GAPDH was used as a loading control. For IF staining, nuclei were counterstained with DRAQ5. Scale bar, 25 µm. Data are presented as mean ± SD. P values were calculated using an unpaired Student’s t-test. *p<0.05; **p<0.01. Quantification of staining and immunoblotting was performed using ImageJ (NIH).

These inhibitory effects of ECM1 on HGF signaling and proliferation induction were further validated in human hepatocyte-derived HepaRG cells. ECM1 suppresses HGF-induced c-MET and ERK phosphorylation (**Figure 4A, B**), MYC nuclear translocation (**Figure 4C**), and expression of cell-cycle regulators (**Figure 4D, E**), resulting in reduced cell proliferation (**Figure 4F, G**). Notably, flow cytometry reveals that HGF promotes G2-phase entry, while ECM1 overexpression causes cell-cycle arrest at the S/G2 transition (**Figure 4H**), in line with its inhibitory effect on BIRC5, CCNA2, and CCNB1 (**Figure 4I**), which are mainly increased during S to G2 progression (Dunajova et al., 2016; Lemmens and Lindqvist, 2019). As expected, forced MYC expression is promoting cell-cycle regulator expression and proliferation in HepaRG cells in the presence of ECM1 overexpression (**Figure 4J-K**).

**Figure 4.**
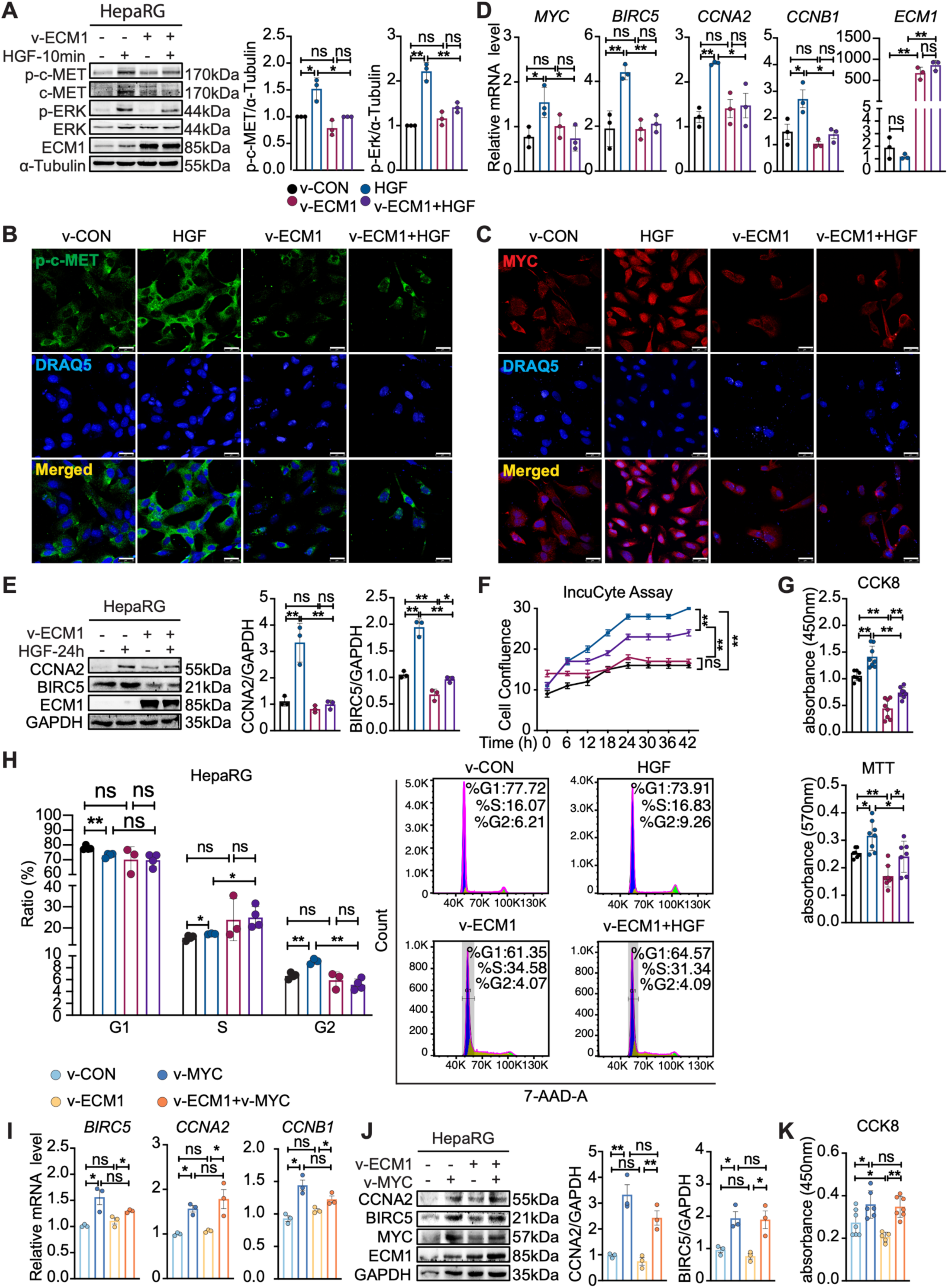
ECM1 overexpression inhibits active HGF-induced HepaRG cell proliferation. **(A, B)** Immunoblotting and IF staining of p-c-MET and p-ERK in HepaRG cells transfected with a control or ECM1 plasmid (v-ECM1) and stimulated with recombinant HGF for 10 min. **(C)** IF staining of MYC in HepaRG cells transfected with control or ECM1 plasmid and treated with HGF for 4 h. **(D, E)** Relative mRNA and protein expression of cell-cycle regulators (BIRC5, CCNA2, CCNB1, CCNB2) in HepaRG cells transfected with control or ECM1 plasmid and treated with HGF for 24 h. **(F, G)** Proliferation of HepaRG cells transfected with control or ECM1 plasmid and treated with HGF for 48 h, assessed by IncuCyte live-cell imaging, MTT cell viability assay, and CCK-8 assay. **(H)** Flow cytometry analysis of HepaRG cells transfected with control or ECM1 plasmid and treated with HGF for 48 h, showing changes in cell-cycle distribution. (7-AAD-A, 7-Aminoactinomycin D-Area) **(I, J)** Relative mRNA and protein expression of cell-cycle genes in HepaRG cells transfected with MYC, ECM1, or both. **(K)** Proliferation of HepaRG cells overexpressing MYC and/or ECM1 by CCK-8 assay. For qRT-PCR, human *PPIA* served as an endogenous control. For immunoblotting, α-Tubulin or GAPDH was used as a loading control. For IF staining, nuclei were counterstained with DRAQ5. Scale bar, 25 µm. Data are presented as mean ± SD. P values were calculated using an unpaired Student’s t-test. *p<0.05; **p<0.01. Quantification of staining and immunoblotting was performed using ImageJ (NIH).

Together, these data demonstrate that ECM1 inhibits HGF-c-Met activation and downstream ERK-MYC signaling, thereby suppressing cell-cycle progression and hepatocyte proliferation.

### ECM1 inhibits HGF-c-MET-ERK-MYC signaling by binding to the active HGF *α*-subunit

We next investigated the mechanism by which ECM1 interferes with active HGF signaling. Using the AlphaFold3 structural prediction system (Abramson et al., 2024), we modeled potential interaction sites between full-length ECM1 and HGF, which suggests direct binding (**Figure 5A**). Co-IF staining of the liver tissue from the mice 2 hours post 70% PHx reveals co-localization of ECM1 and HGF (**Figure 5B**). To validate this interaction, including physiological context, we performed proximity ligation assays (PLA) in liver tissue from healthy mice 2 hours following 70% PHx, a time point characterized by physiological Ecm1 expression and enhanced HGF levels. PLA signals are detectable in the extracellular space of the liver tissue (**Figure 5C**, upper panel). *In vitro*, robust PLA signals are present in PMHs co-expressing Ecm1 and HGF (**Figure 5C**, lower panel). Further, Ecm1 overexpression and recombinant (r)Ecm1 incubation obviously reduce the binding of the HGF and its receptor c-Met, as identified by the PLA (**Figure 5D**). Negative controls using either the primary or PLA probe alone for the PLA experiments are shown in **Figure S4A-C**. To determine whether the observed interaction is direct, we performed *in vitro* pull-down assays using purified recombinant proteins, which confirm that ECM1 directly binds to the HGF α-subunit, which also contains the high-affinity c-MET-binding domain (**Figure 5E**; full immunoblots shown in **Figure S4D**).

**Figure 5.**
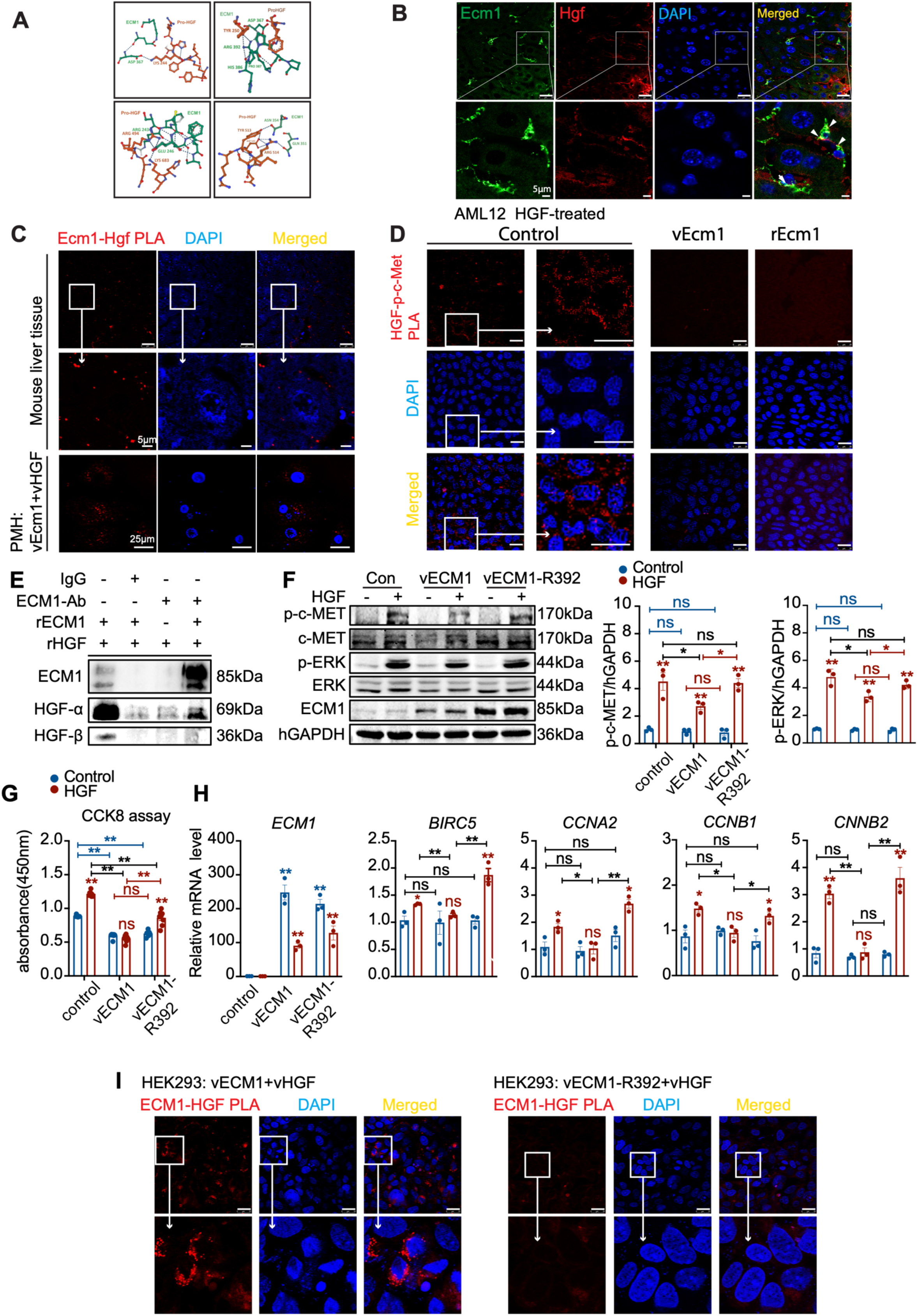
ECM1 directly binds to HGF α-subunit through residue R392. **(A)** Structural prediction of ECM1-HGF interaction generated by AlphaFold3 using full-length input sequences. The ECM1 protein and residues involved in the interaction were shown in green while pro-HGF was shown in red. Models were visualized in Visual Studio Code and edited with Protein Viewer (v0.1.0). **(B)** Co-IF staining and co-localization analyses of ECM1 and HGF in healthy mouse liver tissues. **(C)** PLA detecting Ecm1-Hgf interaction in liver tissues from control mice 2 hours post 70% PHx, and in PMHs co-overexpressing ECM1 and HGF. **(D)** PLA detecting HGF-c-MET interaction in AML12 cells overexpressed with Ecm1 or incubated with recombinant Ecm1. **(E)** *In vitro* pull-down assay of recombinant human ECM1 and HGF proteins. **(F)** Immunoblotting of p-c-MET and p-ERK in HepaRG cells transfected with control, WT ECM1 or ECM1-R392 mutant plasmid and stimulated with recombinant HGF for 10 min. **(G)** Proliferation of HepaRG cells transfected with control, ECM1, or ECM1-R392 mutated plasmid and treated with HGF for 48 h, assessed by CCK-8 assay. **(H)** Relative mRNA expression of *ECM1* and cell-cycle regulators (*BIRC5*, *CCNA2*, *CCNB1*, *CCNB2*) in HepaRG cells transfected with control, ECM1 or ECM1-R392 mutated plasmid and treated with HGF for 24 h. **(I)** PLA detecting ECM1-HGF interaction in HEK293 cells co-overexpressing ECM1 or ECM1-R392 and HGF. For immunoblotting, GAPDH was used as a loading control. For qRT-PCR, mouse *Ppia* was used as an endogenous control. For IF staining, nuclei were counterstained with DAPI. Scale bar, 25 µm. Data are presented as mean ± SD. P values were calculated by an unpaired Student’s t-test. *p<0.05; **p<0.01.

AlphaFold3 predictions and previous structural work (Uchikawa et al., 2021) suggest ECM1 residues R243, D367, and R392 are potential determinants for HGF-binding. We therefore generated three ECM1 mutants harboring alanine substitutions at each residue and expressed them in HepaRG cells prior to HGF stimulation. In contrast to WT ECM1, the ECM1-R392 mutant fails to suppress HGF-induced c-MET and ERK phosphorylation and shows a markedly reduced ability to inhibit cell proliferation, as well as mRNA expression of cell-cycle regulatory genes (**Figure 5F-H**), while ECM1-R243/D392 mutation does not show an obvious difference compared to the WT form. Next, to determine whether the R392 mutation disrupts the interaction between ECM1 and HGF, we performed PLA in HEK293 cells co-overexpressing HGF and either WT ECM1 as a positive control, or the ECM1-R392 mutant. Robust PLA signals indicating ECM1-HGF interaction are detected in cells expressing WT ECM1 **(Figure 5I**, left), whereas PLA signals are almost undetectable in cells expressing the ECM1-R392 mutant **(Figure 5I**, right). Negative controls using either the primary antibody or PLA probe alone in HEK293 cells are shown in **Figure S4E, F.** These results indicate that residue R392 is critical for ECM1-HGF binding and for ECM1-mediated inhibition of HGF signaling.

To determine whether ECM1 interferes with active HGF signaling by directly binding to its receptor c-MET, we again used AlphaFold 3 to predict potential interactions based on protein structures (Abramson et al., 2024). No interaction is predicted between ECM1 and the extracellular domain of c-MET. Consistently, *in vitro* pull-down assays of recombinant ECM1 and c-MET proteins reveal no direct binding (**Figure S4G**). Moreover, co-IF staining of Ecm1 and c-Met in wild-type mouse liver tissues 2 hours following operation shows no evidence of co-localization (**Figure S4H**).

Together, these findings demonstrate that ECM1 inhibits HGF signaling by direct binding to the HGF α-chain with a critical role for residue R392, thereby preventing its engagement with c-MET and subsequent downstream activation.

### Proliferative hepatocytes in liver disease patients are localized in ECM1-negative regions

To confirm the translatability of our findings to patients with liver disease and especially those with significant regenerative activity, we co-stained human serial liver tissue sections for ECM1 and proliferation markers. In relatively healthy liver tissue (n=4), ECM1 is abundant in the extracellular space, and the hepatocytes are negative for MYC and PCNA (**Figure 6A**, upper panel). In contrast, cirrhotic livers with regeneration nodules from patients with chronic liver disease (CLD) (n=12) showed almost complete loss of ECM1 expression, whereas hepatocytes proliferated, indicated by nuclear staining for MYC and PCNA (**Figure 6A**, lower panel). We also analysed biopsies from acute liver failure (ALF) patients with either spontaneous recovery (n=4), or those who progressed and received liver transplantation (n=29). In recovered patients, ECM1 expression in the extracellular matrix is largely rescued with surrounding hepatocytes being negative for nuclear MYC and PCNA (**Figure 6B**, upper panel). Notably, a subset of the explanted livers from progressing ALF patients shows the presence of ECM1 expression in some areas (ALF liver transplantation (LT)x-1, n=15), albeit at reduced levels compared with the recovered cohort, whereby nuclear MYC and PCNA staining are consistently lacking in ECM1-positive regions (**Figure 6B**, middle panel). Presence of nuclear MYC and PCNA staining is most prominent in ECM1-negative ALF patients (ALF LTx-2, n=14) (**Figure 6B**, lower panel). Quantification confirms that ECM1 expression negatively correlates with hepatocyte proliferative activity in cirrhosis and ALF groups (Figure **6C****, D**).

**Figure 6.**
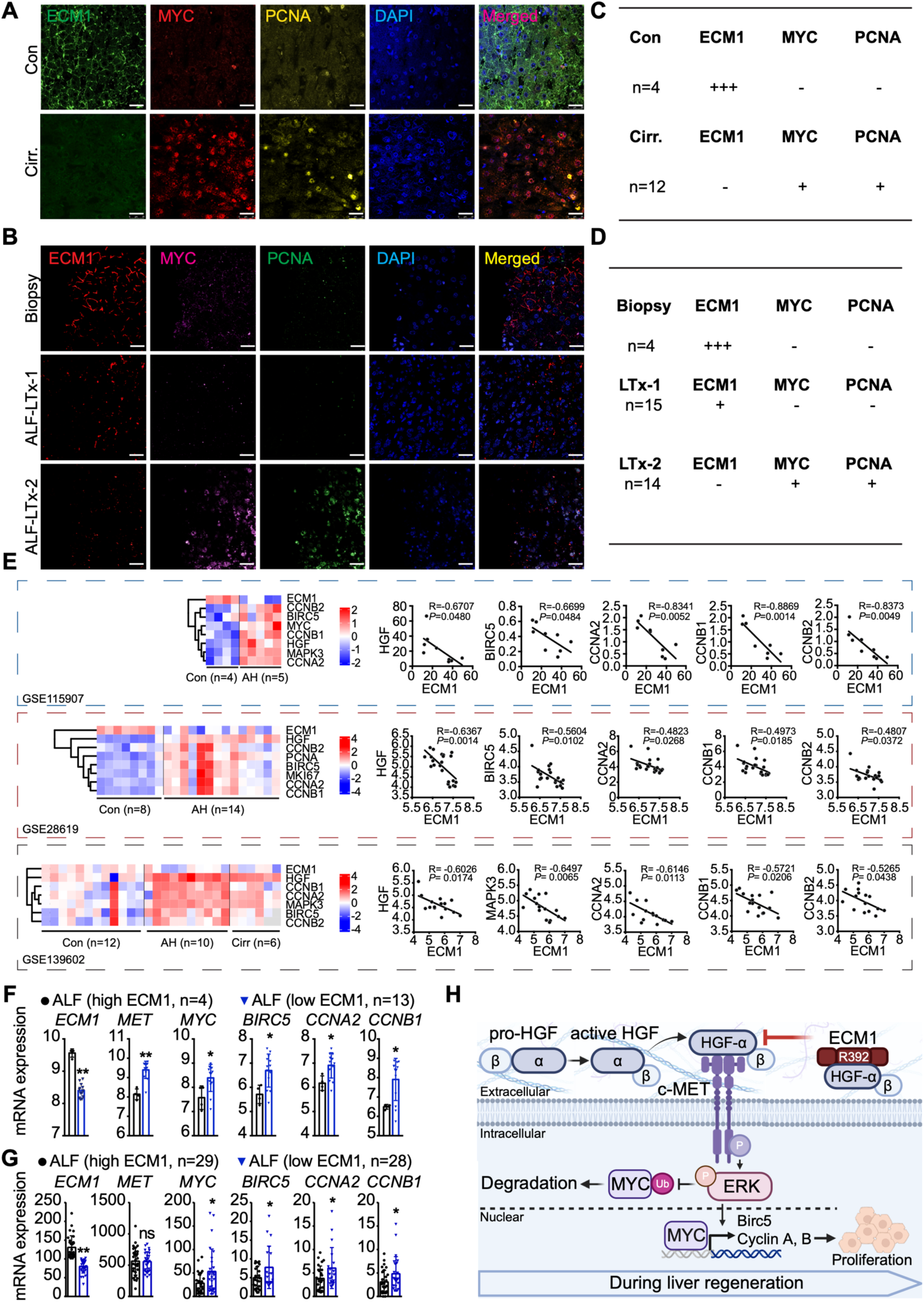
Decreased ECM1 is associated with enhanced proliferative signaling in patient livers. **(A)** Representative IF staining of ECM1, MYC, and PCNA in liver tissues from healthy controls and cirrhotic patients with regenerating nodules. **(B)** Representative IF staining of ECM1, MYC, and PCNA in acute liver failure (ALF) patients who recovered (biopsy samples) or underwent liver transplantation (LTx-1 and LTx-2). **(C, D)** Quantification of ECM1, MYC, and PCNA staining in control, cirrhotic, and ALF liver tissues. **(E)** Heatmap and correlation analyses of ECM1 and HGF pathway target genes using publicly available datasets (GSE155907, GSE28619, GSE139602). **(F, G)** Differential mRNA expression of ECM1 and HGF target genes in ALF patient cohorts (GSE38941, GSE208413). **(H)** Schematic model showing ECM1-mediated inhibition of HGF-c-MET-ERK-MYC signaling and hepatocyte proliferation through direct binding to the HGF α-subunit, a process critically dependent on ECM1 residue R392. For IF staining, nuclei were counterstained with DAPI. Scale bar, 25 µm. Data are presented as mean ± SD. P values were 5calculated using an unpaired Student’s t-test. *p<0.05; **p<0.01.

To extend these observations, we analysed publicly available patient RNA-seq datasets from liver disease patients, including alcoholic hepatitis (AH; GSE155907, GSE28619) (Affo et al., 2013), Acute-on-chronic liver failure (ACLF) (GSE139602) (Graupera et al., 2022), and ALF (GSE38941, GSE208413) (Nissim et al., 2012; Starlinger et al., 2023). Across all cohorts, *ECM1* expression is significantly reduced relative to healthy controls, whereas HGF signaling components, including *HGF*, *MAPK3*, *MYC*, *BIRC5*, *CCNA2*, *CCNB1*, and *CCNB2*, are markedly upregulated (**Figure 6E**). Correlation analyses reveal that ECM1 expression consistently shows an inverse relationship with HGF, its signaling components and its downstream targets involved in cell proliferation across multiple datasets (GSE155907, GSE28619, GSE139602) (**Figure 6E**). Stratifying the ALF cohorts (GSE38941, GSE208413) along ECM1 expression levels, the ECM1-low group exhibits significantly higher expression of the HGF target genes (**Figure 6F, G**). Collectively, the clinical analyses demonstrate that loss of ECM1 is robustly associated with enhanced HGF signaling and proliferative activity in hepatocytes, reinforcing our experimental findings that ECM1 functions as a negative regulator of liver regeneration.

In conclusion, we propose the following model: Upon pro-HGF proteolytically cleaved into α and β subunits, with the α-subunit harboring the high-affinity c-Met-binding domain, homeostatic ECM1 levels sequester the HGF α-chain and prevent c-MET engagement, with ECM1 residue R392 playing an essential role in this interaction. Thus, rapid downregulation of ECM1 is necessary to initiate liver regeneration along the HGF-c-Met axis, which involves ERK phosphorylation, MYC nuclear translocation, and induction of cell-cycle activation, leading to hepatocyte proliferation and an increase in liver mass (**Figure 6H**).

### HSC-secreted ECM1-HGF regulates hepatocyte proliferation after PHx

To identify the liver cell population responsible for the rapid downregulation of ECM1 early after PHx and to examine cell-cell communication associated with the ECM1-HGF axis during regeneration, we analysed publicly available single-cell RNA sequencing (scRNA-seq) dataset (GS151309), compromising liver tissues from adult mice and mice at 24, 48, and 96 hours after 70% PHx (Chembazhi et al., 2021). After quality control filtering, 270,209 cells are retained for downstream analysis. Following batch correction, principal component analysis, graph based clustering, and UMAP embedding, 26 transcriptionally distinct clusters are identified (**Figure S5A**). Marker based annotation classifies these clusters into eight major cell populations, including hepatocytes, quiescent hepatic stellate cells (qHSCs), activated hepatic stellate cells (aHSCs), liver sinusoidal endothelial cells (LSECs), NK and T cells, B cells, dendritic cells, and neutrophils (**Figure S5B**). Hepatocytes accounts for the largest proportion of cells across all time points. Among non-parenchymal cells, LSECs represents the most abundant population and shows a transient increase at 24 hours after PHx.

Average expression analysis shows that Ecm1 and Hgf are most highly expressed in qHSCs among the annotated cell types (**Figure S5C, D**). Time course analysis shows that Ecm1 expression in qHSCs decreases at 24 hours after PHx, remains low at 48 hours, and increases again at 96 hours (**Figure 7A, B**). In contrast, Hgf expression shows higher levels during the early phase after PHx, particularly in qHSCs and LSECs (**Figure 7B**). Hepatocytes are identified as the primary responders to HGF signaling and the main proliferative cell population during regeneration (**Figure S5E, F**). Consistently, activation of the HGF-c-MET-ERK-MYC signaling cascade in hepatocytes is inversely correlated with *Ecm1* expression in qHSCs (**Figure 7A, C**) , as gene set variation analyses (GSVA) of hepatocytes shows increased enrichment scores for HGF-c-Met signaling, RAS/MAPK, cell cycle related gene sets, and KEGG cell cycle. Further, reduced *Ecm1* expression in qHSCs is associated with increased hepatocyte proliferation, as identified by the Mki67 expression levels (**Figure 7D**). Collectively, these analyses suggest that, after PHx, qHSCs coordinate a shift toward reduced ECM1 and increased HGF production, thereby enhancing HGF availability to activate c-MET signaling in hepatocytes and promote proliferative liver mass recovery (**Figure 7E**).

**Figure 7.**
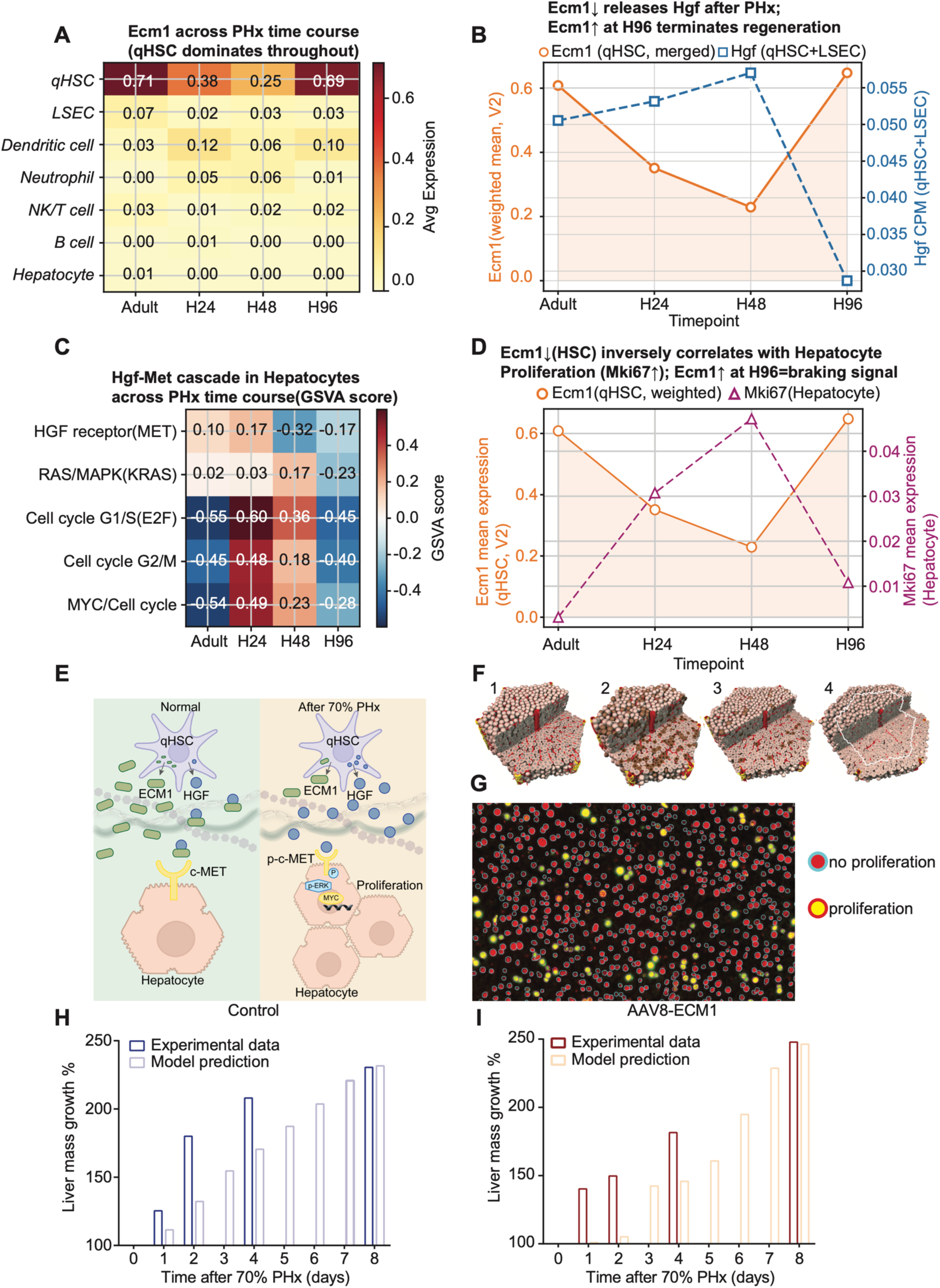
HSC-secreted ECM1-HGF regulates hepatocyte proliferation after PHx. **(A)** Heatmap showing Ecm1 expression across annotated cell types in adult mouse liver and at 24, 48, and 96 hours after partial hepatectomy. **(B)** Time course plot showing changes in Ecm1 and Hgf expression after partial hepatectomy. **(C)** Heatmap of GSVA enrichment scores for HGF MET related pathways in hepatocytes across the regeneration time course, including MET signaling, PI3K AKT mTOR, mTORC1, STAT3, RAS MAPK, immediate early gene response, MYC and cell cycle programs, E2F G1S, G2M, and KEGG cell cycle. **(D)** Time course plot showing Ecm1 expression in quiescent hepatic stellate cells and Mki67 expression in hepatocytes after partial hepatectomy. **(E)** Scheme showing ECM1-HGF-mediated crosstalk between HSCs and hepatocytes after 70% PHx. **(F)** Visualization of 3D model simulation (lobule scale, control scenario). Colors: Red: Lobular vessels and sinusoids, yellow: bile ducts, brown cells: proliferating. **(1)** t=0 days after PHx, **(2)** t=2 days, **(3)** t=4 days **(4)** t=8 days. **(G)** Visualization of automated segmentation of liver tissue by TiQuant/LiKiSeg. (Colors indicate detection of cell proliferation state: Cyan outline around red nuclei: no proliferation, red outline around yellow nuclei: proliferation). **(H)** Plot: Normalized liver mass change after PHx for C57BL/6J mice (control) and **(I)** AAV8-ECM1-treated mice for the experiment and model.

To further study the role of ECM1 in the orchestration of structural and functional changes during initiation, maintenance, and termination of liver regeneration across multiple length scales, and the contribution of ECM1 to cell-cell crosstalk, we utilized a detailed three-dimensional spatio-temporal multiscale (3D-STM) model of liver regeneration after 70% PHx that integrates key structural and functional complexity of this process (Hoehme et al., 2023). Model simulations captured the full experimental time frame of 8 days after surgery (PHx or sham) for control and AAV8-ECM1 groups. We directly compared experimental and simulated data of cell proliferation rates derived from BrdU-staining at 0, 2, 4 and 8 days after surgery (**Figure 7F**), supported by automated tissue segmentation using the established TiQuant/LikiSeg software tools (Hoehme et al., 2023) (**Figure 7G**), as well as liver mass recovery dynamics (Hoehme et al., 2023) (**Figure 7H, I**). Quantification of the corresponding model simulations clearly shows that the experimental data for both cohorts are fully consistent with a mainly HGF/c-Met controlled proliferation response to 70% PHx extended by the ECM1-dependent regulation of HGF signalling (**Figure 7G-I**). Notably, the experimental data obtained from AAV8-ECM1-treated mice cannot be explained without this ECM1-based regulation of proliferation, or a functionally equivalent regulatory mechanism.

Additionally, the model predicted proliferation rates at time points not experimentally measured, allowing verification of its underlying assumptions. For example, at day 3 post-PHx, the model predicts a proliferation rate of approximately 0.64 per day in the AAV8-ECM1 cohort, representing a ∼3.6-fold increase relative to controls (**Figure 7H, I**). At later time points (days 5-7), the model predicts an increase in proliferation (∼0.22 per day on average), approximately 2.4-fold higher than in controls (**Figure 7H, I**). These significantly increased proliferation rates are predicted to be required for the experimentally measured catch-up recovery of liver mass, which even slightly exceeds that of the control group on day 8 (**Figure 7H, I**). Consequently, a future experimental measurement of cell proliferation at the predicted time points will provide evidence for the validity of the model. Collectively, this integrative modelling framework supports the experimental findings and provides a quantitative platform to test mechanistic hypotheses and guide future experiments refining ECM1-mediated regulation of liver regeneration.

## Discussion

Liver regeneration is a highly orchestrated process that integrates mitogenic signaling, extracellular matrix (ECM) remodeling, and dynamic cytokine networks (Fausto, 2000; Michalopoulos, 2007; Michalopoulos and Bhushan, 2021). Here, we identify the extracellular matrix component ECM1 as a negative regulator of hepatocyte proliferation in this process. Initially, we found that ECM1 expression is rapidly downregulated following injury, such as acute carbon tetrachloride (CCl4) treatment (Hammad et al., 2023), and in our case, PHx. In order provide support if the observed ECM1 depletion is functionally required for regeneration, we rescued the downregulation by enforced ECM1 expression using AAV8-ECM1 injection before the PHx. This leads to impaired HGF-c-MET-ERK-MYC signaling, suppressed cell-cycle progression, and delayed liver mass recovery. Mechanistically, ECM1 binds the HGF α-chain and its residue R392 playing a pivotal role, preventing HGF interaction with c-Met. Consistent with this inhibitory effect, hepatocytes in ECM1-negative regions of patients with cirrhosis (regeneration nodules area) or acute liver failure exhibit robust proliferative signals. Additionally, liver disease patient omics datasets consistently show a correlation of ECM1 downregulation with HGF pathway activation and cell cycle progression. Together, these findings uncover ECM1 as a molecular checkpoint that restrains hepatocyte proliferation during regeneration in acutely or chronically damaged livers. Additionally, we found that the ECM1 overexpression reduces total HGF protein levels in the liver tissue (**Figure 2B**). One possible mechanism is that the HGF α-chain bound by ECM1 is targeted for protein degradation, a hypothesis that requires further investigation to elucidate the pathways governing HGF stability and turnover.

Our study also indicates variant and context-dependent functions of ECM1 as a scaffold matrisomal protein that especially orchestrates activation of proforms of ECM-deposited cytokines. We previously reported that epidermal growth factor (EGF) signaling mediates homeostatic ECM1 expression in healthy liver, which is necessary to suppress spontaneous latent TGF-β activation and avoid fibrogenesis (Li et al., 2025). Mechanistically, ECM1 inhibits latent TGF-β activation by interfering with its activators, including integrin, ADAMTS1, thrombospondin-1, MMP-2/9 (Fan et al., 2019; Link et al., 2025). In chronic disease settings, ECM1 thus serves a protective role by preventing TGF-β-driven fibrosis, and thereby represents a promising drug target. By contrast, our current findings reveal an opposite requirement for the initiation of liver regeneration upon acute injury. Here, transient downregulation of ECM1 is required to relieve its inhibitory effect on HGF, thereby permitting c-MET-ERK-MYC activation and proliferative expansion of hepatocytes. This regulatory switch ensures that hepatocytes can efficiently re-enter the cell cycle in response to acute injury. Once sufficient regenerative hepatocytes are generated, ECM1 expression must be restored to terminate proliferative signaling, thereby preventing uncontrolled growth, tumorigenesis, or progression to hepatocellular carcinoma (HCC).

Proper termination of liver regeneration requires tightly regulated inhibitory signals, among which TGF-β is a well-established terminating suppressor of hepatocyte proliferation (Houck and Michalopoulos, 1989) TGF-β expression increases early after PHx and stays elevated until the end of regeneration (Braun et al., 1988). Upon expression, TGF-β is secreted as a large and inactive complex, and deposited in the extracellular matrix, and from which the active cytokine will be generated by conformational change or proteolytic cleavage of latency associated peptide by integrin or MMPs (Li et al., 2022). Our previous work demonstrated that ECM1 inhibits latent TGF-β activation, suggesting that restoration of ECM1 expression during the later phase of regeneration may contribute to interference with the TGF-β mediated termination of the proliferative response. This mechanism may partly explain why liver-to-body weight ratios in AAV8-ECM1-treated mice eventually catch up to those of control animals despite an early delay in regeneration. Furthermore, liver regeneration requires dynamic remodeling of the extracellular matrix, which involves a temporary surge of TGF-β signaling to reorganize tissue architecture and restore homeostasis (Karkampouna et al., 2012). However, excessive or sustained TGF-β release risks chronic activation of HSCs, driving regeneration towards fibrosis. At this stage, ECM1 re-emerges as a crucial brake, limiting redundant TGF-β activity to safeguard tissue repair without tipping toward excessive apoptosis or fibrogenesis. Taken together, these observations underscore the plasticity of ECM1 function across distinct injury contexts. ECM1 operates as a molecular switch: suppressing TGF-β in chronic injury to prevent fibrosis, but transiently downregulates in acute regeneration to permit HGF-driven proliferation, before returning to maintain homeostasis and prevent pathological remodeling. We propose that ECM1 functions as a “Hepatostat” in the regenerating liver, a specific “sensor molecule” that regulates growth-related signals (e.g, HGF and TGF-β) so that the liver size is properly maintained (**Figure S6**).

Given that EGF is besides HGF also an important regulator of LR (Fausto et al., 1995; Michalopoulos and Bhushan, 2021), and we previously found that EGF signaling mediates homeostatic ECM1 expression in healthy hepatocytes (Li et al., 2025), it needs further investigation to explore the interaction between ECM1 and EGF in the context of liver regeneration. According to previous studies, expression of a hepatocyte-specific dominant-negative EGFR delays liver regeneration, but the process proceeds to completion, due to a compensatory enhanced c-MET pathway activation (Lopez-Luque et al., 2016). In contrast, HGF/c-MET signaling is not redundant and necessary for liver regeneration, as mice depleted of c-Met in hepatocytes are hypersensitive to apoptosis, display persistent inflammation and exhibit impaired recovery (Huh et al., 2004). Therefore, in this study, we focused on the interaction between ECM1 and HGF and its effect on liver regeneration. Nevertheless, the mechanism of EGF signaling-mediated transcriptional regulation of ECM1 expression and its role in coordinating regeneration versus fibrosis warrants further investigation.

Our findings also raise intriguing translational implications. In acute liver failure or extensive resection, premature or sustained ECM1 expression could be detrimental by inhibiting HGF-driven regeneration. Conversely, in chronic fibrotic settings, restoring ECM1 expression may be beneficial for restraining aberrant latent TGF-β activation and fibrosis progression. Future studies are warranted to dissect how upstream regulators, including HGF, EGF, and other factors, differentially modulate ECM1 expression across injury contexts, and how therapeutic manipulation of ECM1 could be leveraged to promote regeneration while limiting fibrosis.

In summary, we demonstrate that ECM1 directly binds the HGF α-chain, thereby inhibiting c-MET-ERK-MYC activation and hepatocyte proliferation during LR. Together with prior work linking ECM1 to HGF/EGF- and TGF-β-mediated pathways, our study identifies ECM1 as a context-dependent regulator of liver repair, serving protective anti-fibrotic roles in chronic injury but functioning as a brake on hepatocyte proliferation in acute regeneration.

## Materials and Methods

Human samples, mouse experiments, chemical reagents, primers, and antibodies used in this study, and further detailed information on the methods are presented in the supplementary materials.

## Competing interests

The authors have declared that no competing financial interests exists.

## Financial support

This work was supported by the Deutsche Forschungsgemeinschaft (DFG) [grant number DO 373/20-1 SD; DO 373/20-2 SD SW], Federal Ministry of Education and Research (BMBF) Program LiSyM-HCC, [grant number PTJ-031L0257A]; to SD, the Stiftung Biomedizinische Alkohol-Forschung, [grant number 73000350], and HiChol [01GM1904A to RL].

## Author contributions

Conceptualization: Sai Wang, Steven Dooley

Methodology: Ye Yao, Yujia Li, Chenjun Huang, Jiayi Ma, Liang Xu, Stephanie D. Wolf, Julian Gilljam, Haifeng Zeng, Stefan Munker, Xioachun Cao-Ehlker, Ruochan Li, Bernhard Renz, Hanno Nieß, Seddik Hammad, Elisa Holstein, Laura Danielczyk, Shanshan Wang, Weiguo Fan, Stefan Höhme, Sai Wang

Investigation: Ye Yao, Yujia Li, Chenjun Huang, Jiayi Ma, Liang Xu, Haifeng Zeng, Laura Danielczyk, Shanshan Wang, Stefan Höhme, Sai Wang

Visualization: Ye Yao, Yujia Li, Chenjun Huang, Jiayi Ma, Liang Xu, Haifeng Zeng, Laura Danielczyk, Shanshan Wang, Stefan Höhme, Sai Wang

Funding acquisition: Sai Wang, Steven Dooley

Project administration: Sai Wang, Steven Dooley

Supervision: Sai Wang, Steven Dooley

Writing - Original: Sai Wang, Steven Dooley, Ye Yao, Stefan Höhme, Peter ten Dijke, Weiguo Fan

Writing - review & editing: Ye Yao, Yujia Li, Chenjun Huang, Jiayi Ma, Liang Xu, Stephanie D. Wolf, Julian Gilljam, Haifeng Zeng, Stefan Munker, Xioachun Cao-Ehlker, Ruochan Li, Bernhard Renz, Hanno Nieß, Seddik Hammad, Elisa Holstein, Laura Danielczyk, Roman Liebe, Honglei Weng, Ursula Klingmüller, Matthias P. A. Ebert, Chunfang Gao, Johannes Bode, Peter ten Dijke, Donato Inverso, Shanshan Wang, Stefan Höhme, Weiguo Fan, Steven Dooley, Sai Wang

## Acknowledgments

We acknowledge the support of Prof. Hellmut Augustin’s group for providing liver tissue samples collected at different time points following 70% partial hepatectomy, which served as an independent experimental cohort. We also acknowledge the support of the LIMa Live Cell Imaging at Microscopy Core Facility Platform Mannheim (CFPM).

## Notes

### Competing Interest Statement

The authors have declared no competing interest.

## References

Abramson, J., Adler, J., Dunger, J., Evans, R., Green, T., Pritzel, A., Ronneberger, O., Willmore, L., Ballard, A.J., Bambrick, J., et al. (2024). Accurate structure prediction of biomolecular interactions with AlphaFold 3. Nature 630, 493–500.

Affo, S., Dominguez, M., Lozano, J.J., Sancho-Bru, P., Rodrigo-Torres, D., Morales-Ibanez, O., Moreno, M., Millan, C., Loaeza-del-Castillo, A., Altamirano, J., et al. (2013). Transcriptome analysis identifies TNF superfamily receptors as potential therapeutic targets in alcoholic hepatitis. Gut 62, 452–460.

Braun, L., Mead, J.E., Panzica, M., Mikumo, R., Bell, G.I., and Fausto, N. (1988). Transforming growth factor beta mRNA increases during liver regeneration: a possible paracrine mechanism of growth regulation. Proc Natl Acad Sci U S A 85, 1539–1543.

Chembazhi, U.V., Bangru, S., Hernaez, M., and Kalsotra, A. (2021). Cellular plasticity balances the metabolic and proliferation dynamics of a regenerating liver. Genome Res 31, 576–591.

Dunajova, L., Cash, E., Markus, R., Rochette, S., Townley, A.R., and Wheatley, S.P. (2016). The N-terminus of survivin is a mitochondrial-targeting sequence and Src regulator. J Cell Sci 129, 2707–2712.

Fajardo-Puerta, A.B., Mato Prado, M., Frampton, A.E., and Jiao, L.R. (2016). Gene of the month: HGF. J Clin Pathol 69, 575–579.

Fan, W., Liu, T., Chen, W., Hammad, S., Longerich, T., Hausser, I., Fu, Y., Li, N., He, Y., Liu, C., et al. (2019). ECM1 Prevents Activation of Transforming Growth Factor beta, Hepatic Stellate Cells, and Fibrogenesis in Mice. Gastroenterology 157, 1352–1367 e1313.

Fausto, N. (2000). Liver regeneration. J Hepatol 32, 19–31.

Fausto, N., Laird, A.D., and Webber, E.M. (1995). Liver regeneration. 2. Role of growth factors and cytokines in hepatic regeneration. FASEB J *9*, 1527-1536.

Gene Ontology, C. (2021). The Gene Ontology resource: enriching a GOld mine. Nucleic Acids Res 49, D325–D334.

Graupera, I., Isus, L., Coll, M., Pose, E., Diaz, A., Vallverdu, J., Rubio-Tomas, T., Martinez-Sanchez, C., Huelin, P., Llopis, M., et al. (2022). Molecular characterization of chronic liver disease dynamics: From liver fibrosis to acute-on-chronic liver failure. JHEP Rep 4, 100482.

Hammad, S., Ogris, C., Othman, A., Erdoesi, P., Schmidt-Heck, W., Biermayer, I., Helm, B., Gao, Y., Pioronska, W., Holland, C.H., et al. (2023). Tolerance of repeated toxic injuries of murine livers is associated with steatosis and inflammation. Cell Death Dis 14, 414.

Hoehme, S., Hammad, S., Boettger, J., Begher-Tibbe, B., Bucur, P., Vibert, E., Gebhardt, R., Hengstler, J.G., and Drasdo, D. (2023). Digital twin demonstrates significance of biomechanical growth control in liver regeneration after partial hepatectomy. iScience 26, 105714.

Houck, K.A., and Michalopoulos, G.K. (1989). Altered responses of regenerating hepatocytes to norepinephrine and transforming growth factor type beta. J Cell Physiol 141, 503–509.

Huh, C.G., Factor, V.M., Sanchez, A., Uchida, K., Conner, E.A., and Thorgeirsson, S.S. (2004). Hepatocyte growth factor/c-met signaling pathway is required for efficient liver regeneration and repair. Proc Natl Acad Sci U S A 101, 4477–4482.

Karkampouna, S., Goumans, M.J., Ten Dijke, P., Dooley, S., and Kruithof-de Julio, M. (2016). Inhibition of TGFbeta type I receptor activity facilitates liver regeneration upon acute CCl4 intoxication in mice. Arch Toxicol 90, 347–357.

Karkampouna, S., Ten Dijke, P., Dooley, S., and Julio, M.K. (2012). TGFbeta signaling in liver regeneration. Curr Pharm Des 18, 4103–4113.

Lemmens, B., and Lindqvist, A. (2019). DNA replication and mitotic entry: A brake model for cell cycle progression. J Cell Biol 218, 3892–3902.

Li, Y., Fan, W., Link, F., Wang, S., and Dooley, S. (2022). Transforming growth factor beta latency: A mechanism of cytokine storage and signalling regulation in liver homeostasis and disease. JHEP Rep 4, 100397.

Li, Y., Huang, C., Fan, W., Hammad, S., Geraud, C., Berger, L., Wang, S., Yao, Y., Tong, C., Rubie, C., et al. (2025). ECM1 expression in chronic liver disease: Regulation by EGF/STAT1 and IFNgamma/NRF2 signalling. JHEP Rep 7, 101423.

Link, F., Li, Y., Zhao, J., Munker, S., Fan, W., Nwosu, Z.C., Yao, Y., Wang, S., Huang, C., Liebe, R., et al. (2025). ECM1 attenuates hepatic fibrosis by interfering with mediators of latent TGF-beta1 activation. Gut 74, 424–439.

Lopez-Luque, J., Caballero-Diaz, D., Martinez-Palacian, A., Roncero, C., Moreno-Caceres, J., Garcia-Bravo, M., Grueso, E., Fernandez, A., Crosas-Molist, E., Garcia-Alvaro, M., et al. (2016). Dissecting the role of epidermal growth factor receptor catalytic activity during liver regeneration and hepatocarcinogenesis. Hepatology 63, 604–619.

Michalopoulos, G. (1993). HGF and liver regeneration. Gastroenterol Jpn 28 *Suppl 4*, 36–39; discussion 53-36.

Michalopoulos, G.K. (2007). Liver regeneration. J Cell Physiol 213, 286–300.

Michalopoulos, G.K. (2010). Liver regeneration after partial hepatectomy: critical analysis of mechanistic dilemmas. Am J Pathol 176, 2–13.

Michalopoulos, G.K. (2011). Liver regeneration: alternative epithelial pathways. Int J Biochem Cell Biol 43, 173–179.

Michalopoulos, G.K., and Bhushan, B. (2021). Liver regeneration: biological and pathological mechanisms and implications. Nat Rev Gastroenterol Hepatol 18, 40–55.

Mizuno, K., Tanoue, Y., Okano, I., Harano, T., Takada, K., and Nakamura, T. (1994). Purification and characterization of hepatocyte growth factor (HGF)-converting enzyme: activation of pro-HGF. Biochem Biophys Res Commun 198, 1161–1169.

Naka, D., Ishii, T., Yoshiyama, Y., Miyazawa, K., Hara, H., Hishida, T., and Kidamura, N. (1992). Activation of hepatocyte growth factor by proteolytic conversion of a single chain form to a heterodimer. J Biol Chem 267, 20114–20119.

Nissim, O., Melis, M., Diaz, G., Kleiner, D.E., Tice, A., Fantola, G., Zamboni, F., Mishra, L., and Farci, P. (2012). Liver regeneration signature in hepatitis B virus (HBV)-associated acute liver failure identified by gene expression profiling. PLoS One 7, e49611.

Roselli, H.T., Su, M., Washington, K., Kerins, D.M., Vaughan, D.E., and Russell, W.E. (1998). Liver regeneration is transiently impaired in urokinase-deficient mice. Am J Physiol 275, G1472–1479.

Starlinger, P., Brunnthaler, L., McCabe, C., Pereyra, D., Santol, J., Steadman, J., Hackl, M., Skalicky, S., Hackl, H., Gronauer, R., et al. (2023). Transcriptomic landscapes of effective and failed liver regeneration in humans. JHEP Rep 5, 100683.

Uchikawa, E., Chen, Z., Xiao, G.Y., Zhang, X., and Bai, X.C. (2021). Structural basis of the activation of c-MET receptor. Nat Commun 12, 4074.

