## Supplementary Tables 1 and 2; Supplementary Figures 1-6. for "Transient suppression of the ECM1 gatekeeper is essential for HGF/c-MET-driven liver regeneration"

**Supplementary information**

Table S1-S2

Supplementary materials and methods

Supplementary Figure S1-S6

References

### Temporal downregulation of ECM1 gatekeeper is required for HGF/c-MET-mediated liver repair

Ye Yao [1#], Yujia Li [1#], Chenjun Huang [1, 2], Jiayi Ma [3], Liang Xu [4], Stephanie D. Wolf [5], Julian Gilljam [5], Haifeng Zeng [1], Stefan Munker [6, 7], Xioachun Cao-Ehlker [6, 7], Ruochan Li [6, 7], Bernhard Renz [6, 7], Hanno Nieß [6, 7], Seddik Hammad [1], Elisa Holstein [8], Laura Danielczyk [1], Roman Liebe [9], Honglei Weng [1], Ursula Klingmüller [8], Matthias P. A. Ebert [1, 10, 11], Chunfang Gao [2], Johannes Bode [5], Donato Inverso[12], Peter ten Dijke [13], Shanshan Wang [3], Stefan Höhme [14], Weiguo Fan [15], Steven Dooley [1*], Sai Wang [1*]

[1] Department of Medicine II, University Medical Center Mannheim, Medical Faculty Mannheim, Heidelberg University, Mannheim, Germany; [2] Department of Clinical Laboratory Medicine Center, Yueyang Hospital of Integrated Traditional Chinese and Western Medicine, Shanghai University of Traditional Chinese Medicine, Shanghai, China; [3] Beijing Institute of Hepatology, Beijing You'an Hospital, Capital Medical University, Beijing, China; [4] Goethe University Hospital, Frankfurt, Germany; [5] Department of Gastroenterology, Hepatology and Infectious Disease, Faculty of Medicine & Düsseldorf University Hospital, Heinrich Heine University, Düsseldorf, Germany; [6] Department of Medicine II, University Hospital, LMU; Munich, Germany; [7] Liver Center Munich, University Hospital, LMU, Munich, Germany; [8] DKFZ, German Cancer Research Center, Heidelberg, Germany; [9] Clinic of Gastroenterology, Hepatology and Infectious Diseases, Otto-von-Guericke-University, Magdeburg, Germany; [10] Molecular Medicine Partnership Unit, European Molecular Biology Laboratory, Heidelberg, Germany; [11] DKFZ-Hector Cancer Institute at the University Medical Center, Mannheim, Germany; [12] Vascular Pathobiology Unit, San Raffaele Scientific Institute, Medical Faculty, Vita-Salute San Raffaele University, Milan, Italy; [13] Oncode Institute and Department of Cell and Chemical Biology, Leiden University Medical Center, Leiden, Netherlands; [14] Interdisciplinary Centre for Bioinformatics, University of Leipzig, Leipzig, Germany. [15] Key Laboratory of Multi-Cell Systems, Shanghai Institute of Biochemistry and Cell Biology, Center for Excellence in Molecular Cell Science, Chinese Academy of Sciences, Shanghai, China.

#### Supplementary Tables

Table S1: Primers for qRT-PCR and constructs

| Primer | Forward | Reverse |
|---|---|---|
| hACTA2 | GTGTTGCCCCTGAAGAGCAT | GCTGGGACATTGAAAGTCTCA |
| hBIRC5 | CGTCCGGTTGCGCTTTCCTTTCTG | GATGGCACGGCGCACTTTCTCC |
| hCCNA2 | CTCTACACAGTCACGGGACAAAG | CTGTGGTGCTTTGAGGTAGGTC |
| hCCNB1 | GACCTGTGTCAGGCTTTCTCTG | GGTATTTTGGTCTGACTGCTTGC |
| hCCNB2 | CAACCAGAGCAGCACAAGTAGC | GGAGCCAACTTTTCCATCTGTAC |
| hECM1 | TGAACCAAATCTGCCTTCCTAAC | GCTGGACTGTGGTAGGTTCCA |
| hMMP-2 | TACAGGATCATTGGCTACACACC | GGTCACATCGCTCCAGACT |
| hMMP-9 | TGTACCGCTATGGTTACACTCG | GGCAGGGACAGTTGCTTCT |
| hMYC | GGCTCCTGGCAAAAGGTCA | CTGCGTAGTTGTGCTGATGT |
| hPPIA | AGGGTTCCTGCTTTCACAGA | CAGGACCCGTATGCTTTAGG |
| hTGFB1 | AGGGCTACCATGCCAACTTC | CCACGTAGTAGACGATGGGC |
| mActa2 | GTCCCAGACATCAGGGAGTAA | TCGGATACTTCAGCGTCAGGA |
| mBirc5 | CTACCGAGAACGAGCCTGATT | AGCCTTCCAATTCCTTAAAGCAG |
| mCCNA2 | TTGTAGGCACGGCTGCTATGCT | GGTGCTCCATTCTCAGAACCTG |
| mCCNB1 | AGAGGTGGAACTTGCTGAGCCT | GCACATCCAGATGTTTCCATCGG |
| mCCNB2 | GCACTACCATCCTTCTCAGGTG | TGTGCTGCATGACTTCCAGGAC |
| mcFos | CGGGTTTCAACGCCGACTA | TTGGCACTAGAGACGGACAGA |
| mcJun | CCTTCTACGACGATGCCCTC | GGTTCAAGGTCATGCTCTGTTT |
| mEcm1 | GCCAGCTCTGTGGAAGTGGA | CCGGAATCTGTTTATGCTTGC |
| mHgf | ACTTCTGCCGGTCCTGTTG | CCCCTGTTCCTGATACACCT |
| mIL6 | TAGTCCTTCCTACCCCAATTTCC | TTGGTCCTTAGCCACTCCTTC |
| mKlkb1 | TTCAACCGAGTGGGTTATTTTGT | TACTGGGCATCTGGGGTGTAG |
| mMmp-2 | CAAGTTCCCCGGCGATGTC | TTCTGGTCAAGGTCACCTGTC |
| mMmp-9 | GGACCCGAAGCGGACATTG | CGTCGTCGAAATGGGCATCT |
| mMyc | AGCCCCTAGTGCTGCATGA | TCCACAGACACCACATCAATTTC |
| mPlau | GCGCCTTGGTGGTGAAAAAC | TTGTAGGACACGCATACACCT |
| mPlaur | AGGACTACCGTGCTTCGGGAAT | ACACGGTCTCTGTCAGGCTGAT |
| mPpia | GAGCTGTTTGCAGACAAAGTT | CCCTGGCACATGAATCCTGG |
| mTGFa | CAGGCTCTGGAGAACAGCACAT | GACACATGCTGGCTTCTCTTCC |
| mTNFa | CCCTCACACTCAGATCATCTTCT | GCTACGACGTGGGCTACAG |
| mTNFR1 | GTGTGGCTGTAAGGAGAACCAG | CACACGGTGTTCTGAGTCTCCT |
| ECM1-R243A | AGCAATGAGCgcgTTCTGTGAGGCC | TCCTCCCACACAAGTTTG |
| ECM1-D367A | GGATACCCTTgcgAAATACTGTGACC | TCCCAGGCCTTCCATG |
| ECM1-R392A | CAGCCCTACTgcgGATGAGTGCTTTGC | GGAGGGTGGCGGCAA |

Table S2. Antibody information

| Antibody | Company | Cat. No. | Dilution |
|---|---|---|---|
| Anti-human ECM1 | abcam | ab126629 | 1:1000 for WB<br>1:200 for IHC/IF |
| Akt | Cell signaling technology | 9272S | 1:1000 |
| Alexa Flour 488 goat anti-rabbit IgG | Invitrogen | A-11008 | 1:200 |
| Alexa Fluor 555 goat anti-mouse IgG | Invitrogen | A-21422 | 1:200 |
| anti-BrdU | BIO-RAD | MCA-6144 | 1:500 |
| HRP anti-His Tag antibody | Biolegend | 652503 | 1:3000 |
| Anti-mouse ECM1 | abcam | ab253158 | 1:1000 for WB<br>1:200 for IHC/IF |
| Birc5 | Cell signaling technology | 71G4B7 | 1:1000 |
| c-Jun | Cell signaling technology | 60A8 | 1:1000 |
| Ccna2 | Cell signaling technology | BF683 | 1:1000 |
| DRAQ5 | Cell signaling technology | 4084L | 1:1000 |
| EGFR | Cell signaling technology | 2232 | 1:1000 |
| Erk | Santa Cruz Biotechnology | sc-1359 | 1:500 |
| GAPDH | abcam | sc-32233 | 1:1000 |
| HGF | R&D | AF-294-NA | 1:1000 for WB |
| HRP-linked anti-mouse | Santa Cruz Biotechnology | sc-2005 | 1:5000 |
| HRP-linked anti-rabbit | Santa Cruz Biotechnology | sc-2357 | 1:5000 |
| c-Met | Cell signaling technology | 25H2 | 1:1000 |
| Myc | Santa Cruz Biotechnology | sc-40 | 1:100 for IHC |
| Myc | abcam | ab32072 | 1:1000 for wb |
| p-Akt | Cell signaling technology | 4060T | 1:1000 |
| p-c-Met | Invitrogen | 44-888G | 1:1000 for wb<br>1:100 for IF |
| p-EGFR | Cell signaling technology | Y1068 | 1:1000 |
| p-Erk | Santa Cruz Biotechnology | sc-7383 | 1:500 |
| p-Stat3 | Cell signaling technology | 9145T | 1:1000 |
| PCNA | abcam | ab252848 | 1:100 for IF |
| PCNA | proteintech | 10205-2-AP | 1:1000 for IF |
| Stat3 | Santa Cruz Biotechnology | sc-482 | 1:200 |
| α-Tubulin | abcam | ab4074 | 1:1000 |

#### Supplementary Materials and Methods

##### Patient samples

The study protocol was approved by the appropriate local ethics committees (LL-2023-027-K). Written informed consent was obtained from patients or their representatives. Relatively healthy liver tissues were obtained by liver biopsy. Liver tissues from patients with cirrhosis or ALF were collected at the time of liver transplantation. Organ allocation and transplantation timing were governed by the China Liver Transplant Registry (CLTR) (Wan et al., 2016), an official scientific registry authorized by the Chinese Health Ministry, and were determined according to individual Model for End-Stage Liver Disease (MELD) scores.

##### Animal studies

Male C57BL/6J mice (8 to 10 weeks old) were purchased from the Javier Lab (Badalona, Spain). The generation and characterization of *Ecm1*-knockout (KO) mice have been described previously (Fan et al., 2019). The *Ecm1*-tdTomato reporter mice were generated by CRISPR-Cas9-mediated targeted integration of an *Ecm1*-tdTomato cassette at the 5' end of the endogenous *Ecm1* locus, as reported in our previous study (Li et al., 2025). *Ecm1*-tdTomato mice received a single tail vein injection of AAV8-ECM1 or AAV8-Control 7 days prior to sham operation or 70% partial hepatectomy (PHx). Liver samples were collected at 2 h, 1 day, 2 days, 4 days, and 8 days post-surgery. Each mouse received 100 µl of AAV containing 1.25 x $10^{11}$ vector genomes. AAV8-Control and AAV8-ECM1 were purchased from VectorBuilder. 70% PHx was performed as previously described (Mitchell and Willenbring, 2008). Sham-operation mice underwent a midline abdominal skin and muscle incision, followed by closure of the peritoneum and skin with sutures, as described previously (Wolf et al., 2023). All experiments were performed using 8-10-week-old male or female mice, with six mice per experimental group. All animal procedures were conducted in accordance with institutional and governmental guidelines for animal care and were approved by the local animal care committee (Regierungspräsidium Karlsruhe, Abteilung 3 - Landwirtschaft, Ländlicher Raum, Veterinär- und Lebensmittelwesen; approval numbers 35-9185.81/G-144/22, G-103/23).

##### Primary hepatocyte isolation

Primary hepatocytes were isolated from 8 -16 weeks old male C57BL/6J mice as previously described (Godoy et al., 2013). Mice were anesthetized by intraperitoneal injection of ketamine hydrochloride (5mg/100mg body weight; 10% solution) and xylazine hydrochloride (1mg/100mg body weight; 2% solution). The liver was sequentially perfused via the inferior vena cava, with simultaneous transection of the portal vein, using 50 ml of perfusion buffer (Krebs-Henseleit buffer with 0.5mM EDTA, Sigma), followed by 50 ml of collagenase A buffer (Krebs-Henseleit buffer with 0.1mM $CaCl_2$ and 0.4mg/ml collagenase A, Sigma). Following perfusion, the liver was excised and gently dissociated using forceps to release hepatocytes into suspension buffer. The cell suspension was passed through a 100-µm cell strainer and centrifuged at 50 g for 5 min at 4°C. The supernatant was discarded, and the cell pellet was resuspended in Percoll solution (prepared in Hank's buffer) to remove dead cells, followed by centrifugation at 50 g for 10 min at 4°C. The resulting hepatocyte pellet was resuspended in culture medium and seeded for subsequent experiments.

**Cell culture and treatment**

Freshly isolated primary mouse hepatocytes (PMHs) were seeded onto 6- or 12-well plates pre-coated with Rat Tail Collagen I (11179179001, Roche, 250µg/mL in 0.2% acetic acid) and cultured in William's E medium (A1217601, ThermoFisher) supplemented with 10% fetal bovine serum (FBS), 2mM L-glutamine, 1% penicillin (100U/mL)-streptomycin (100µg/mL) P/S, 40 ng/ml dexamethasone, and 0.5% Insulin-Transferrin-Selenium (ITS). HepaRG cells were maintained in William's E medium supplemented with 10% FBS, 1% L-glutamine, 1% P/S, 50µM hydrocortisone hemisuccinate, and 5 µg/ml insulin. For hepatocyte differentiation, HepaRG cells were cultured in differentiation medium consisting of William's E medium supplemented with 10% FBS, 1% L-glutamine, 1% P/S, 1x Insulin-Transferrin-Selenium (ITS), 40ng/ml dexamethasone, and 10mM nicotinamide. AML12 cells were cultured in DMEM/F-12 medium (21331-020, Gibco) supplemented with 10% FBS, 2mM L-glutamine, 1% P/S, 1x ITS, and 40ng/ml dexamethasone. HEK293T cells were cultured in DMEM medium (11965092, Life Technologies) supplemented with 10% FBS, 1% P/S. All cells were cultured at 37°C in a humidified incubator containing 5% $CO_2$.

After overnight attachment, PMHs, HepaRG, AML12, and HEK293 cells were serum-starved for 4-6 h prior to transfection with mouse or human ECM1 expression plasmids using Lipofectamine® reagents (L3000015, Invitrogen life technologies, USA) according to the manufacturer's instructions. 24 or 48 h after transfection, cells were treated with HGF (50 ng/ml) or pro-EGF/EGF (20/40 ng/ml) for 10 min, 24 h, or 48 h, as indicated.

**RNA isolation and real-time quantitative (RT-q) PCR**

All reagents and consumables used for RNA extraction were RNase-free or treated with 0.1% diethyl pyrocarbonate (DEPC; 4387937, Thermo Fisher Scientific) to eliminate RNase contamination. Total RNA was isolated from liver tissues or cultured cells using TRIzol (15596018, Invitrogen) according to the manufacturer's instructions. For cDNA synthesis, 500 ng of total RNA was reverse-transcribed using random primers (SO142, Thermo Fisher Scientific), RevertAid H Minus Reverse Transcriptase (EP0452, Thermo Fisher Scientific), and RiboLock RNase Inhibitor (EO0382, Thermo Fisher Scientific). qRT-PCR was performed on a StepOnePlus Real-Time PCR system (Applied Biosystem) using SYBR Green Master Mix. Relative gene expression levels were normalized to the housekeeping gene *Ppia*/*PPIA*. Three biological replicates were analysed for each condition. Relative fold changes in target gene expression were calculated using the $-2^{\Delta\Delta CT}$ formula (Livak and Schmittgen, 2001). Primer sequences for qRT-PCR were listed in **Supplementary Table 1**.

**Generation of mutant ECM1 plasmid**

Three different point mutations (R243A, D367A, R392A) in the ECM1 plasmid were mutagenized using a Q5 Site-Directed Mutagenesis Kit (New England Biolabs, E0554S) according to the manufacturer's instructions. Briefly, the protocol has three steps, including exponential amplification (PCR), kinase, ligase & DpnI (KLD) treatment, and transformation, all constructed plasmid sequences were verified by Sanger sequencing (Eurofins). The used mutagenic primers were listed in **Supplementary Table 1**.

**Immunoblotting**

Liver tissues or cultured cells were lysed on ice for 10 min in RIPA buffer (1% Triton X-100, 50mM Tris [pH 7.5], 300mM NaCl, 1mM EGTA, 1mM EDTA and 0.1% SDS) supplemented with protease inhibitors (36978, Thermo Fisher Scientific) and phosphatase inhibitors (P5726, Sigma-Aldrich). Lysates were clarified by centrifugation at 13,000 rpm for 15min at 4°C, and supernatants were collected. Protein concentrations were determined using the Bio-Rad protein assay kit, with absorbance measured at 562 nm using a Tecan Infinite M200 plate reader. Equal amounts of protein (30 μg) were mixed with 4× LDS sample buffer (NP0007, Life Technologies), boiled at 99°C for 10 min, and resolved by SDS–PAGE using 8–12% gels. Proteins were transferred onto 0.2-μm nitrocellulose membranes (10600001, GE Healthcare Life Sciences). Membranes were blocked with 5% (w/v) skim milk in TBST (Tris-buffered saline containing 0.05% Tween-20) for 1 h at room temperature, followed by incubation with primary antibodies overnight at 4°C. After washing with TBST, membranes were incubated with appropriate HRP-conjugated secondary antibodies for 1 h at room temperature. Immunoreactive bands were visualized using Western Lightning Plus ECL (NEL103001EA, PerkinElmer) and detected with a Fusion SL4 imaging system (PEQLAB, Germany). A list of antibodies used is provided in **Supplementary Table 2**. Each immunoblotting experiment was independently repeated at least three times.

#### ELISA

HGF concentrations were measured using a human HGF ELISA kit (R&D Systems, DHG00B) according to the manufacturer's instructions. Tissues were lysed, homogenized, and clarified by centrifugation, and the resulting supernatants were analyzed. Standards and samples were added to microplates pre-coated with an HGF-specific capture antibody. After incubation and washing, an HRP-conjugated detection antibody and substrate solution were applied. The reaction was terminated with the provided stop solution, absorbance was measured at 450 nm, and HGF concentrations were calculated from a standard curve.

#### Proximity ligation assay

Proximity ligation assay (PLA) was performed using Duolink® In Situ PLA fluorescence reagents (Sigma-Aldrich) according to the manufacturer's instructions. MPH cultured on coverslips and paraffin-embedded mouse liver sections were deparaffinized, rehydrated, and subjected to antigen retrieval. Samples were permeabilized, blocked, and incubated with rabbit anti-ECM1 (ab253158) and goat anti-HGF (AF-294-NA) primary antibodies diluted in Duolink® antibody diluent. Species-specific PLA probes (PLUS and MINUS) were applied, followed by ligation and amplification. Negative controls included samples incubated with only one primary antibody or with PLA probes alone. Fluorescent signals were detected according to the kit protocol, imaged by Leica confocal microscopy.

#### Immunofluorescence (IF) staining

Fresh liver tissues were embedded in optimal cutting temperature (O.C.T.) compound (TTEK, Hartenstein), snap-frozen on dry ice, and stored at -80°C. Cryosections (6 μm thick) were prepared using a cryostat. For cell-based assays, cultured cells were seeded onto 12-well plates containing glass coverslips (0111580, MARIENFELD). Tissue sections and cultured cells were fixed with 4% paraformaldehyde (PFA) for 15 min at room temperature, followed by three washes with PBS. Samples were permeabilized and blocked with PBS containing 0.5% Triton X-100 and 1% bovine serum albumin (BSA) for 1 h at room temperature. Slides were then incubated with primary antibodies diluted in PBS containing 1% BSA overnight at 4°C. The following day, samples were washed three times with PBS and incubated with fluorophore-conjugated secondary antibodies and DRAQ5 nuclear stain diluted in PBS for 1 h at room temperature in the dark. After three additional PBS washes, slides were mounted using

Fluoromount-G mounting medium (00-4958-02, Invitrogen). Images were acquired using a Leica TCS SP8 confocal microscope. Antibodies used for immunofluorescence are listed in **Supplementary Table 2**.

#### Immunohistochemistry

Paraffin-embedded human liver tissue sections or tissue microarrays (4 μm thick) were deparaffinized and rehydrated using standard protocols. For hematoxylin and eosin (H&E) staining, sections were stained with hematoxylin to visualize nuclei, followed by eosin staining to label cytoplasmic components. For immunohistochemical staining with antibodies, antigen retrieval was performed by boiling sections for 15 min in citrate buffer (pH 6.0) or Tris–EDTA buffer (pH 9.0). After cooling at room temperature for 30 min, endogenous peroxidase activity was quenched by incubation with blocking peroxide solution (S200389-2; Dako) for 45 min. Sections were then washed with PBS and incubated with primary antibodies overnight at 4°C. Following PBS washes, sections were incubated with appropriate secondary antibodies for 1 h at room temperature. Signal detection was performed using 3,3′-diaminobenzidine (DAB), followed by hematoxylin counterstaining. Sections were subsequently dehydrated and mounted using Malinol mounting medium (C9368; Sigma-Aldrich). Slides were scanned using a Leica DMRBE microscope at 20× magnification. A list of antibodies used is provided in **Supplementary Table 2**.

#### *In vitro* pull-down assay

Recombinant ECM1 (1 μg) and/or active HGF (1 μg) proteins were incubated with either control IgG or anti-ECM1 antibody (1 μg) in 200 μL RIPA buffer supplemented with protease and phosphatase inhibitor cocktail (1:100 dilution) at 4°C for 8 h with gentle rotation. Subsequently, 200 μL of protein A/G agarose slurry was added to each reaction, and the mixture was incubated overnight at 4°C. Protein-antibody complexes were collected by centrifugation at 2,000 rpm for 5 min, and supernatants were discarded. The agarose beads were washed three times with PBS containing protease and phosphatase inhibitors (1:1000 dilution). After the final wash, precipitated proteins were resuspended in 2× SDS loading buffer at a volume equivalent to that of the protein A/G agarose beads and boiled for 5 min prior to SDS-PAGE analysis. For input controls, 10 ng of recombinant ECM1 and HGF proteins were loaded directly onto the gel.

#### Single-cell RNAseq analyses

The scRNA-seq dataset (GSE151309), including liver tissue from control group and mice that received 70% PHx, collected at 1, 2, or 4 days after the operation, was analyzed. After quality control filtering, cells with 200 to 8000 detected genes and less than 30% mitochondrial reads were retained for downstream analysis. Batch effects across samples were corrected using BEER. Principal component analysis was performed, and the top 20 principal components were used for dimensionality reduction and clustering. A shared nearest neighbor graph was constructed, and graph based clustering was performed at a resolution of 1.2. Cells were visualized using UMAP embedding. Cell type annotation was performed based on canonical marker gene expression. Hepatocytes were annotated by Hnf4a, Cyp2e1, and Slc27a5. Quiescent hepatic stellate cells were annotated by Lrat, Reln, and Hhip. Activated hepatic stellate cells were annotated by Col1a1 and Lox. Liver sinusoidal endothelial cells were annotated by Stab2, Lyve1, and Oit3. NK and T cells were annotated by Nkg7, Ncr1, and Klrb1c. B cells were annotated by Cd19 and Ms4a1. Dendritic cells were annotated by Siglech and Clec9a. Neutrophils were annotated by Ly6g and Cxcr2. Cell type composition was compared across time points. Average normalized expression of Ecm1 and Hgf was calculated for each annotated cell type. Temporal changes in Ecm1 expression in quiescent hepatic stellate cells and Mki67 expression in hepatocytes were evaluated across the regeneration time course. GSVA was performed in hepatocytes to assess pathway activity related to HGF MET signaling, cell cycle progression, PI3K AKT mTOR, mTORC1, STAT3, RAS MAPK, and immediate early gene response.

**Three-dimensional spatio-temporal multiscale (3D-STM) model**

Since the base model is described in detail in (Hoehme et al., 2023), we will here only reiterate fundamental characteristics of the model, focusing on novel aspects of the model. We use an agent-based model (ABM) approach in which individual cells of liver tissue are explicitly represented as spherical homogeneous visco-elastic objects. For this reason, this type of model is also called an individual-cell-based model, or more recently a digital twin of the liver. In the model, biomechanical interactions between objects, such as hepatocytes or hepatic stellate cells, are modeled using the Johnson-Kendall-Roberts (JKR) contact model. The JKR model includes repulsive and adhesive forces and has been shown to be a valid and robust description of living cells with intact cytoskeletons (Chu et al., 2005). Hepatocyte polarity is taken into account by modifying the adhesive properties of the cell surface based on cell orientation. Cell division is modeled by cell volume growth and concurrent deformation into a dumbbell-like shape, eventually leading to the separation of two independent daughter cells.

Cell-cycle entry is modeled by a detailed intracellular HGF/c-Met signaling model based on ODEs as described in (D'Alessandro et al., 2015). Every model cell simultaneously runs its own instance of this ODE sub-model of intracellular signaling and thereby can independently react to changes in its biomechanical and biochemical microenvironment. The signaling sub-model was extended to include the hypothesized impact of ECM1 up/down-regulation by phenomenologically implementing it as normalized inverse linear scaling prerequisite for HGF-activation. Accordingly, ECM1 is parameterized in both experimental scenarios using mRNA and protein-level analyses.

In addition to this HGF/c-Met signaling-based cell cycle control, biomechanical constraints on cell volume growth are applied, such that for a cell to enter the cell cycle, the pressure in its microenvironment must not exceed a threshold value $p_{div,max}$. Cell movement is a result of force balance summing forces of inertia, friction, JKR-interaction forces and an additional active force for cell migration which includes uniformly distributed random micro motility and directed terms modeling chemotaxis and mechanotaxis. Other cell types like endothelial cells, hepatic stellate cells (HSC) and macrophages are modeled conceptually similar. Sinusoidal blood vessels and larger hepatic veins and arteries are constructed by connected and partially overlapping spherical objects also utilizing the JKR model for interactions with other model objects. In general, all model parameters are derived from experimental data or known physiological values as listed in (Hoehme et al., 2023). Model simulations can be run in partitions of liver tissue of arbitrary size and shape, only limited by computational complexity and memory. Common partitions are individual liver lobules, or cubes of liver tissue with 1mm edge length, typically containing 50-100 lobules.

#### AlphaFold prediction

The predicted structure modelling of HGF and ECM1 were generated using AlphaFold3 (https://alphafoldserver.com/) (Abramson et al., 2024) with full-length of the target proteins as the input sequences. Opened with Visual Studio Code and edited with Protein Viewer V0.1.0.

#### Data availability

The gene expression omnibus (GEO) DataSets GSE166868, GSE155907, GSE28619, GSE139602, GSE38941, and GSE208413 can be accessed via the following link: https://www.ncbi.nlm.nih.gov/gds. All other data are available in the main text or the supplementary materials. Detailed information not provided within the text will be made available upon request via e-mail to either of the corresponding authors.

#### Statistical analysis

Statistical analyses were performed using GraphPad Prism version 6.0. The two-tailed Student's t-test was used to compare two independent groups. A one-way ANOVA was used to test for statistical differences in the means of two groups. Variables were described by mean and standard deviation (SD). Statistical significance was indicated as follows: *$P < 0.05$; **$P < 0.01$. All experiments were repeated at least 3 times independently.

#### Supplementary figures and figure legends

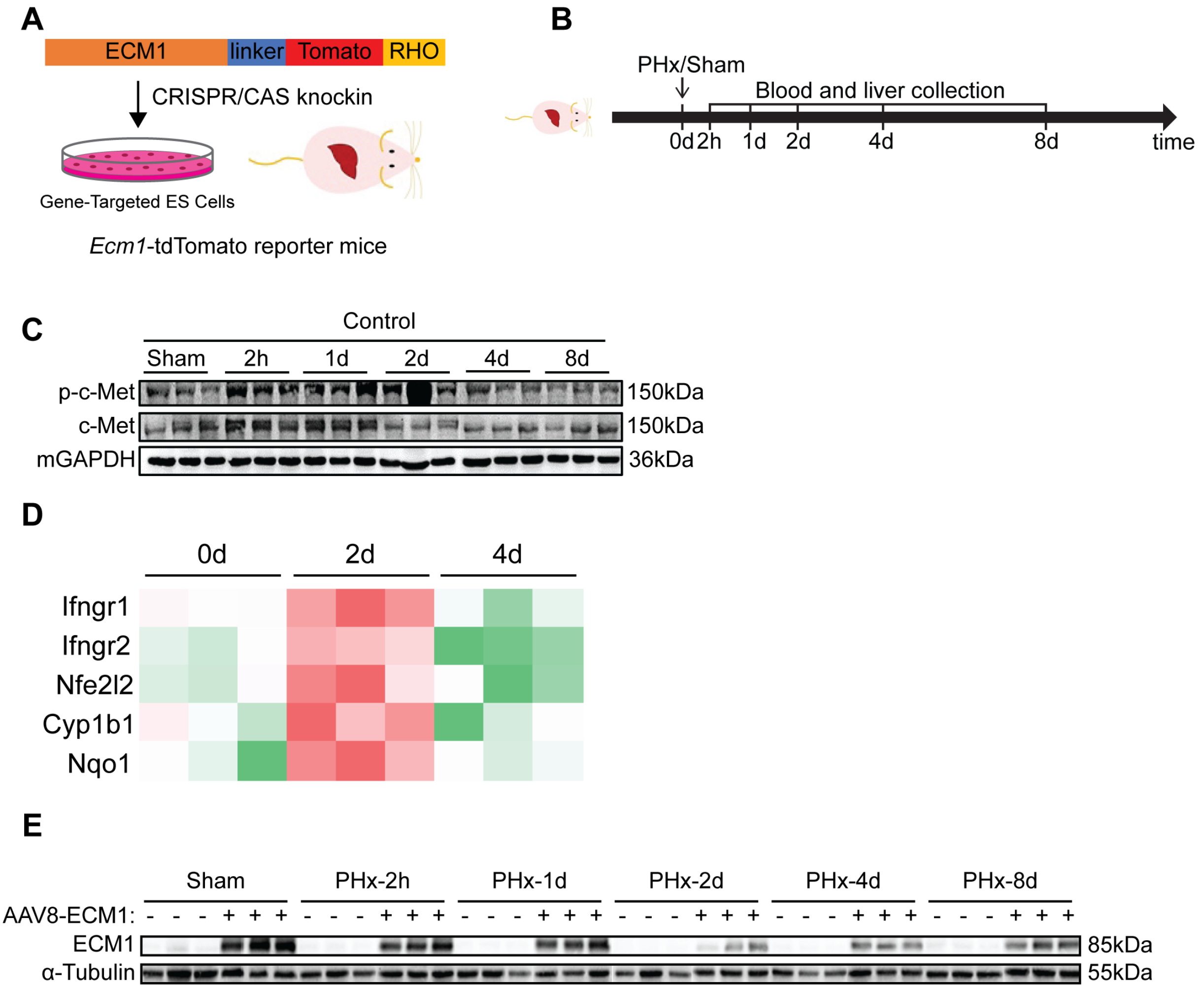


**Figure S1. 70% PHx followed by AAV8-ECM1 intravenous injection in Ecm1-tdTomato mice.**

**(A)** The generation of Ecm1-tdTomato mice using CRISPR/CAS knockin. **(B)** Scheme showing the experimental design of AAV8-ECM1 injection and 70% PHx in Ecm1-tdTomato mice. **(C)** Immunoblotting of p-c-Met and total c-Met in the liver tissue of mice with sham or 70% PHx operation. **(D)** RNA-seq analysis of IFN-γ/NRF2 pathways in liver tissues collected at 2 or 4 days post sham or 70% PHx. **(E)** Immunoblotting of Ecm1 in the liver tissue of mice received 70% PHx with control or AAV8-ECM1 injection. Gapdh/α-Tubulin was used as loading control.

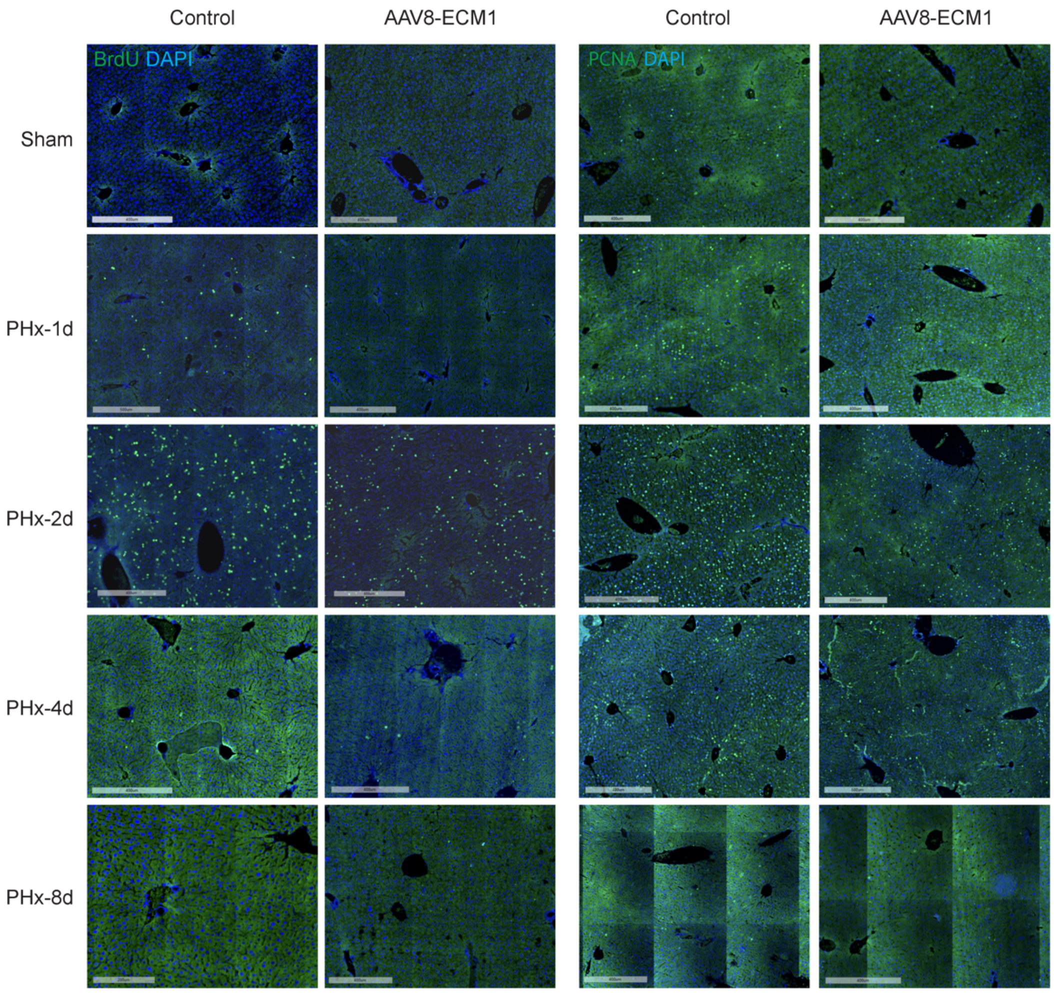


**Figure S2. Proliferation analyses of the mouse liver tissue.**

Representative images of the whole slide scan of the IF staining for BrdU or PCNA in the liver tissue from mice subjected to 70% PHx followed by control or AAV8-ECM1 injection. DAPI was used for nuclear staining. Scale bar, 400 μm.

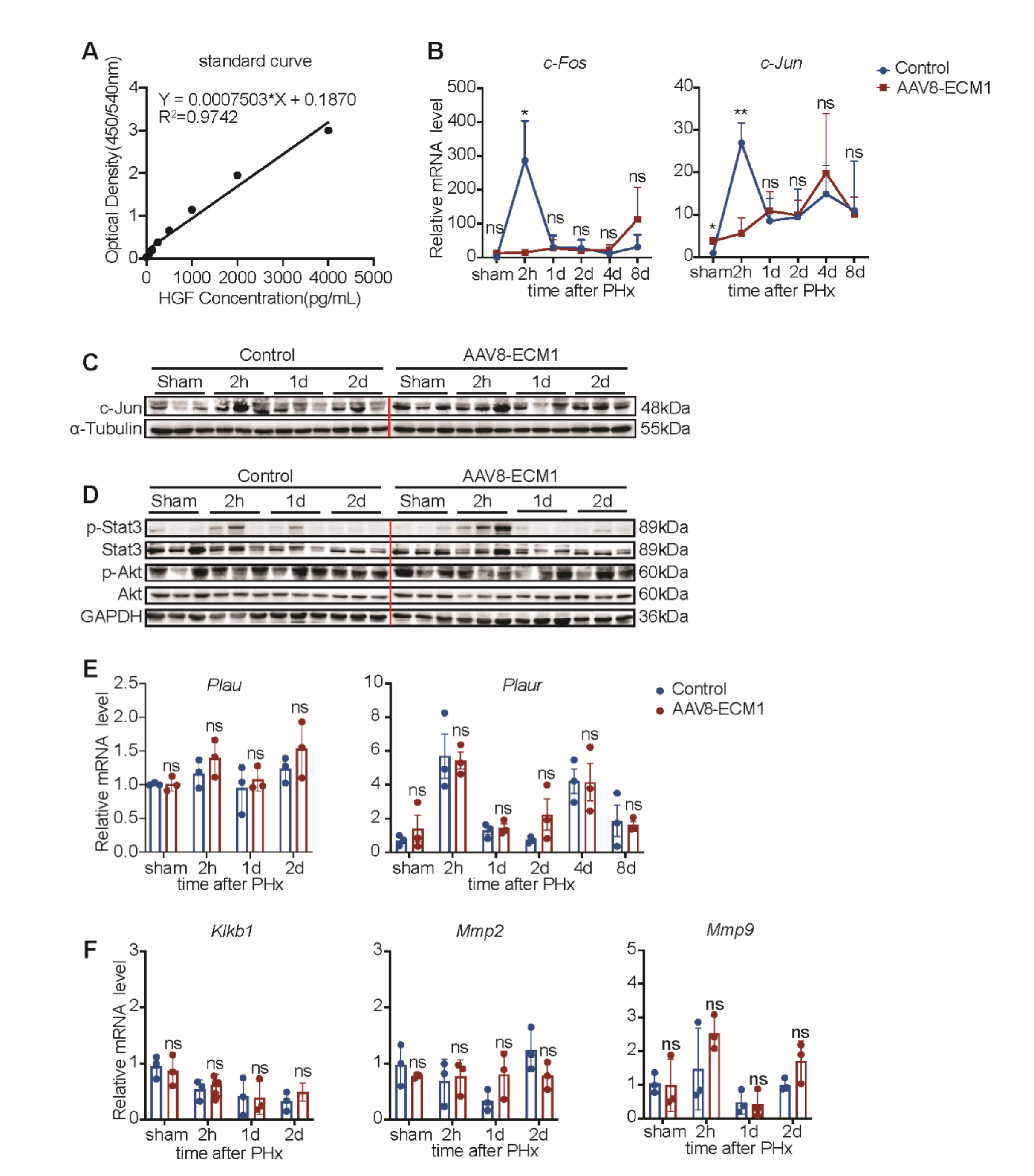


**Figure S3. Analyses of the genes or signaling pathways involved in liver regeneration (LR).**

**(A)** Standard curve of the HGF ELISA assay. **(B)** mRNA expression of *c-Fos* and *c-Jun* in the liver tissue received 70% PHx and/or AAV8-ECM1 injection. **(C, D)** Immunoblotting of c-Jun, p-Stat3, Stat3, p-Akt and Akt in the mouse liver tissue with 70% PHx and/or AAV8-ECM1 injection. **(E, F)** mRNA expression of pro-HGF activators in the liver tissue from mice that received an operation or AAV8-ECM1 injection. For qRT-PCR, mouse *Ppia* was used as endogenous control. For immunoblotting, Gapdh was used as loading control.

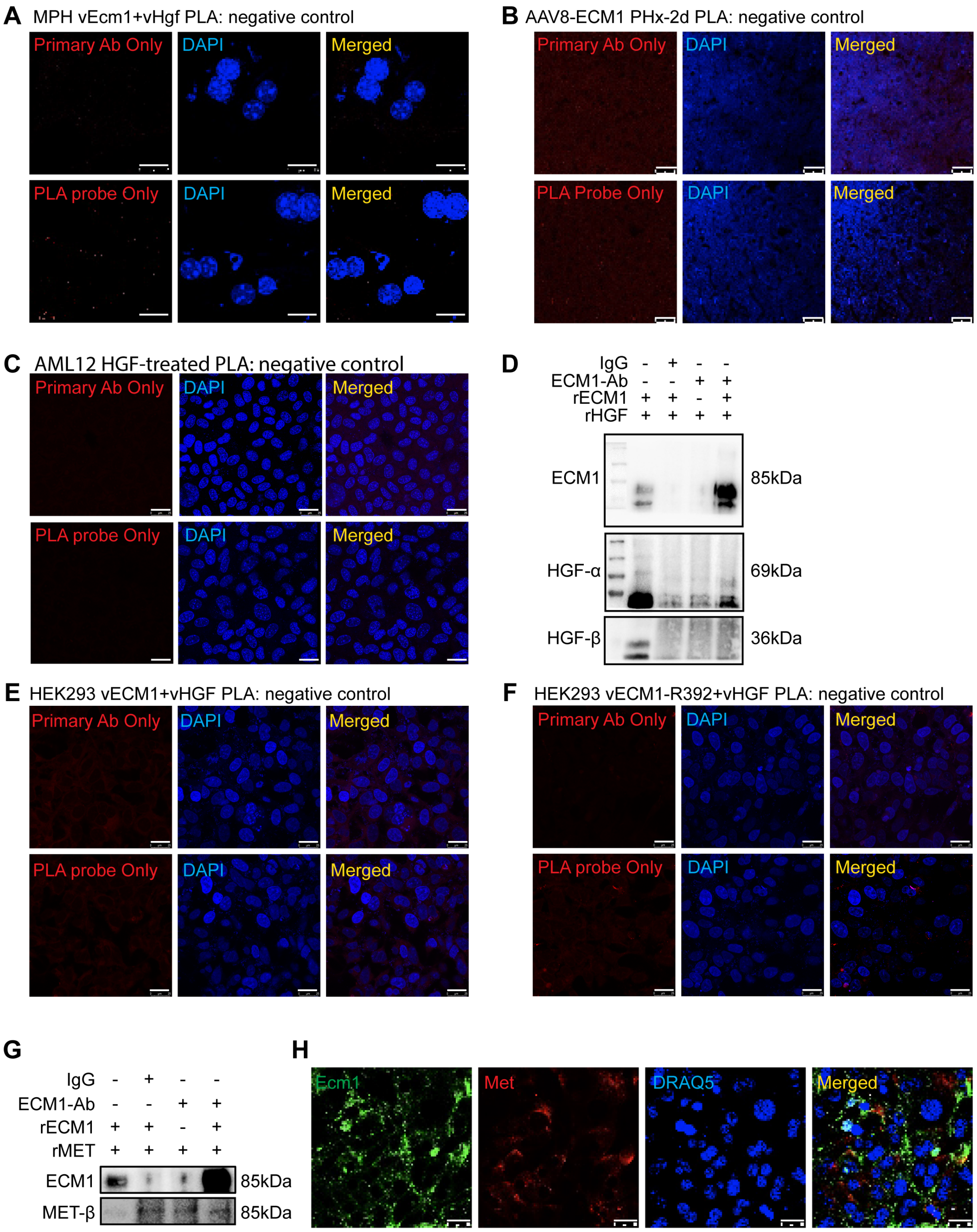


288 **Figure S4. Interaction analyses of ECM1 and its potential binding proteins.**

289 **(A-C)** Negative controls of the PLA assay in the main Figure 6D-F. **(D)** The whole
290 immunoblotting membrane of the *in vitro* pull-down assay of recombinant ECM1 and HGF
291 protein. **(E, F)** Negative controls of the PLA assay in the main Figure 6J. **(G)** *In vitro* pull-down

assay of recombinant ECM1 and c-MET protein. **(H)** Co-IF staining of ECM1 and c-Met in the healthy mouse liver tissue. DAPI or DRAQ5 was used for nuclear staining. Scale bar, 25 μm.

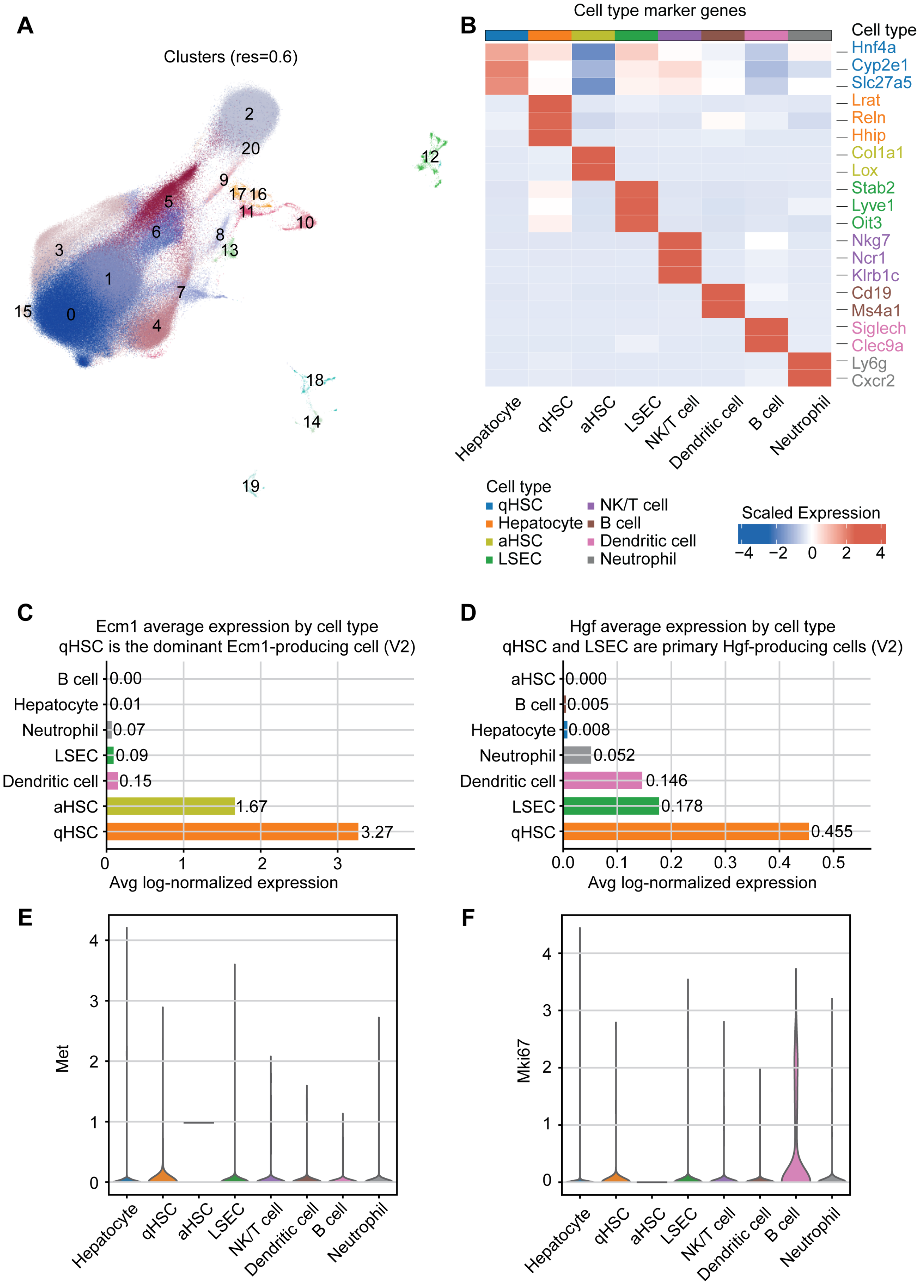

**Figure S5. HSC-secreted ECM1-HGF regulates hepatocyte proliferation after PHx**

**(A)** UMAP visualization of cell clusters from scRNA-Seq (GSE151309). **(B)** Cell type-specific marker genes were utilized for annotating cell populations. **(C-F)** Expression levels of *Ecm1*, *Hgf*, *Met*, and *Ki67* in different cell types.

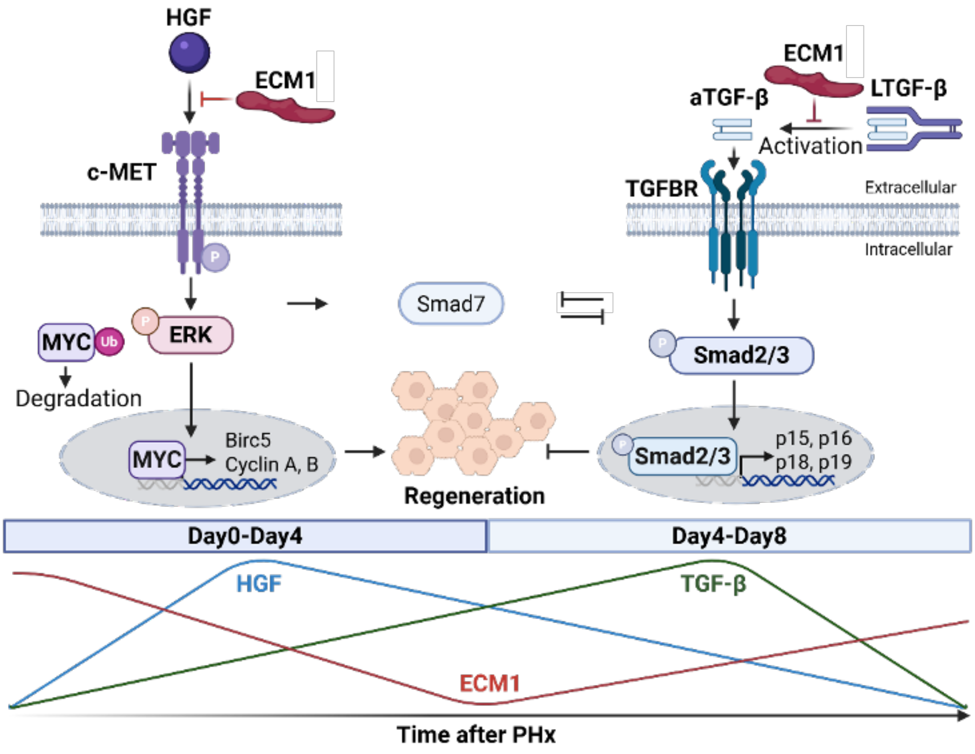


**Figure S6. Schematic representation showing ECM1 as a "hepatostat" coordinating liver size during regeneration.**

At the early phase of LR (day 0-4 post 70% PHx, left panel), rapid downregulation of ECM1 removes an extracellular brake on HGF/c-MET-ERK-MYC signaling, thereby permitting hepatocyte proliferation and initiation of regeneration. At the late stage of LR (day 4-8 post 70% PHx, right panel), restoration of ECM1 expression restrains growth factor signaling and limits TGF-β activation, preventing excessive apoptosis or fibrogenesis and contributing to termination of the regenerative response.

#### References

Abramson, J., Adler, J., Dunger, J., Evans, R., Green, T., Pritzel, A., Ronneberger, O., Willmore, L., Ballard, A.J., Bambrick, J., *et al.* (2024). Accurate structure prediction of biomolecular interactions with AlphaFold 3. Nature *630*, 493-500.

Chu, Y.S., Dufour, S., Thiery, J.P., Perez, E., and Pincet, F. (2005). Johnson-Kendall-Roberts theory applied to living cells. Phys Rev Lett *94*, 028102.

D'Alessandro, L.A., Samaga, R., Maiwald, T., Rho, S.H., Bonefas, S., Raue, A., Iwamoto, N., Kienast, A., Waldow, K., Meyer, R., *et al.* (2015). Disentangling the Complexity of HGF Signaling by Combining Qualitative and Quantitative Modeling. PLoS Comput Biol *11*, e1004192.

Fan, W., Liu, T., Chen, W., Hammad, S., Longerich, T., Hausser, I., Fu, Y., Li, N., He, Y., Liu, C., *et al.* (2019). ECM1 Prevents Activation of Transforming Growth Factor beta, Hepatic Stellate Cells, and Fibrogenesis in Mice. Gastroenterology *157*, 1352-1367 e1313.

Godoy, P., Hewitt, N.J., Albrecht, U., Andersen, M.E., Ansari, N., Bhattacharya, S., Bode, J.G., Bolleyn, J., Borner, C., Bottger, J., *et al.* (2013). Recent advances in 2D and 3D in vitro systems using primary hepatocytes, alternative hepatocyte sources and non-parenchymal liver cells and their use in investigating mechanisms of hepatotoxicity, cell signaling and ADME. Arch Toxicol *87*, 1315-1530.

Hoehme, S., Hammad, S., Boettger, J., Begher-Tibbe, B., Bucur, P., Vibert, E., Gebhardt, R., Hengstler, J.G., and Drasdo, D. (2023). Digital twin demonstrates significance of biomechanical growth control in liver regeneration after partial hepatectomy. iScience *26*, 105714.

Li, Y., Huang, C., Fan, W., Hammad, S., Geraud, C., Berger, L., Wang, S., Yao, Y., Tong, C., Rubie, C., *et al.* (2025). ECM1 expression in chronic liver disease: Regulation by EGF/STAT1 and IFNgamma/NRF2 signalling. JHEP Rep *7*, 101423.

Livak, K.J., and Schmittgen, T.D. (2001). Analysis of relative gene expression data using real-time quantitative PCR and the 2(-Delta Delta C(T)) Method. Methods *25*, 402-408.

Mitchell, C., and Willenbring, H. (2008). A reproducible and well-tolerated method for 2/3 partial hepatectomy in mice. Nat Protoc *3*, 1167-1170.

Wan, P., Xu, D., Zhang, J., Li, Q., Zhang, M., Chen, X., Luo, Y., Shen, C., Han, L., and Xia, Q. (2016). Liver transplantation for biliary atresia: A nationwide investigation from 1996 to 2013 in mainland China. Pediatr Transplant *20*, 1051-1059.

Wolf, S.D., Ehlting, C., Muller-Dott, S., Poschmann, G., Petzsch, P., Lautwein, T., Wang, S., Helm, B., Schilling, M., Saez-Rodriguez, J., *et al.* (2023). Hepatocytes reprogram liver macrophages involving control of TGF-beta activation, influencing liver regeneration and injury. Hepatol Commun *7*.
